# Phylogenetic Permulations: a statistically rigorous approach to measure confidence in associations between phenotypes and genetic elements in a phylogenetic context

**DOI:** 10.1101/2020.10.14.338608

**Authors:** Elysia Saputra, Amanda Kowalczyk, Luisa Cusick, Nathan Clark, Maria Chikina

## Abstract

The wealth of high-quality genomes for numerous species has motivated many investigations into the genetic underpinnings of phenotypes. Comparative genomics methods approach this task by identifying convergent shifts at the genetic level that are associated with traits evolving convergently across independent lineages. However, these methods have complex statistical behaviors that are influenced by non-trivial and oftentimes unknown confounding factors. Consequently, using standard statistical analyses in interpreting the outputs of these methods leads to potentially inaccurate conclusions. Here, we introduce phylogenetic permulations, a novel statistical strategy that combines phylogenetic simulations and permutations to calculate accurate, unbiased p-values from phylogenetic methods. Permulations construct the null expectation for p-values from a given phylogenetic method by empirically generating null phenotypes. Subsequently, empirical p-values that capture the true statistical confidence given the correlation structure in the data are directly calculated based on the empirical null expectation. We examine the performance of permulation methods by analyzing both binary and continuous phenotypes, including marine, subterranean, and long-lived large-bodied mammal phenotypes. Our results reveal that permulations improve the statistical power of phylogenetic analyses and correctly calibrate statements of confidence in rejecting complex null distributions while maintaining or improving the enrichment of known functions related to the phenotype. We also find that permulations refine pathway enrichment analyses by correcting for non-independence in gene ranks. Our results demonstrate that permulations are a powerful tool for improving statistical confidence in the conclusions of phylogenetic analysis when the parametric null is unknown.

## Introduction

Despite the availability of complete genomes for many species, identifying the genetic elements responsible for a phenotype of interest is difficult because there are millions of genetic differences between nearly every species. One strategy to link genotypes and phenotypes is to take advantage of convergent evolutionary events in which multiple unrelated species have evolved similar characteristics. Such events represent natural biological replicates of evolution during which species may have experienced similar genetic changes driving similar phenotypic changes. When lineages independently evolve or lose a shared phenotype, parallel shifts in selective constraints acting on a set of genetic elements indicate that those elements may be involved in the phenotypic shift.

Taking this strategy, several methods have been developed to address the task of linking genetic elements to phenotypes. We have previously developed a method called RERconverge (Kowalczyk et al. 2019; Partha et al. 2019) to link genetic elements to convergently evolving phenotypes based on evolution across a sequence of interest. Our method has been successfully used to identify the genetic basis of adaptation to a marine habitat (Chikina et al. 2016), regression of ocular structures in a subterranean habitat (Partha et al. 2017), and evolution of extreme lifespan and body size phenotypes (Kowalczyk et al. 2020) in mammals. Other groups have developed similar methods that identify the genetic elements underlying a phenotype by identifying similarities in genomes from species that have independently evolved said phenotype. HyPhy RELAX uses sophisticated branch-site models in a likelihood ratio framework to identify genes under relaxation of evolutionary constraint or directed evolution (Wertheim et al. 2015). PhyloAcc calculates substitution rates along a phylogeny in a Bayesian framework to identify convergent rate acceleration in association with phenotypic convergence (Hu et al. 2019). Forward Genomics performs calculations of percent sequence change along a phylogeny to detect similar changes among phenotypically similar species (Hiller et al. 2012). These methods can be broadly categorized as approaches for correlating evolutionary rates (ERs) with species phenotypes and are powerful tools to associate genetic elements with phenotypes. However, our recent work indicates that thorough statistical handling is required when performing analyses on these genomic datasets to avoid both overstating and understating confidence in their results.

### Problems

#### Problem 1: A non-uniform null distribution for p-values from association statistics linking sequence evolution to phenotype evolution

The common approach adopted by all the methods listed above is to use a statistical test to calculate the association between convergent rate acceleration and convergent phenotypic evolution, followed by multiple hypothesis testing corrections to control for Type I error. If an enrichment of small p-values is observed, then it is presumed that some genes (or other genetic elements) are truly associated with the phenotype. This conclusion rests on the assumption that no such relationship existed (i.e., the null hypothesis holds for all genes) and a uniform distribution would be observed. However, our analyses using RERconverge show atypical statistical behavior in which the expected uniform distribution is not observed when null phenotypes are analyzed for gene-phenotype associations (**Figure 1A**). This observation implies that there is considerable data structure (non-independence) among branch-level evolutionary rates that results in correlated test statistics and biased adjusted p-values (Allison et al. 2002; Nettleton et al. 2006; Hu et al. 2010). This problem invalidates statements of significant gene-phenotype correlations that are based on standard multiple hypothesis testing corrections. In other words, a gene may show an apparently significant adjusted p-value in the absence of any true association with the phenotype studied.

**Figure 1.**
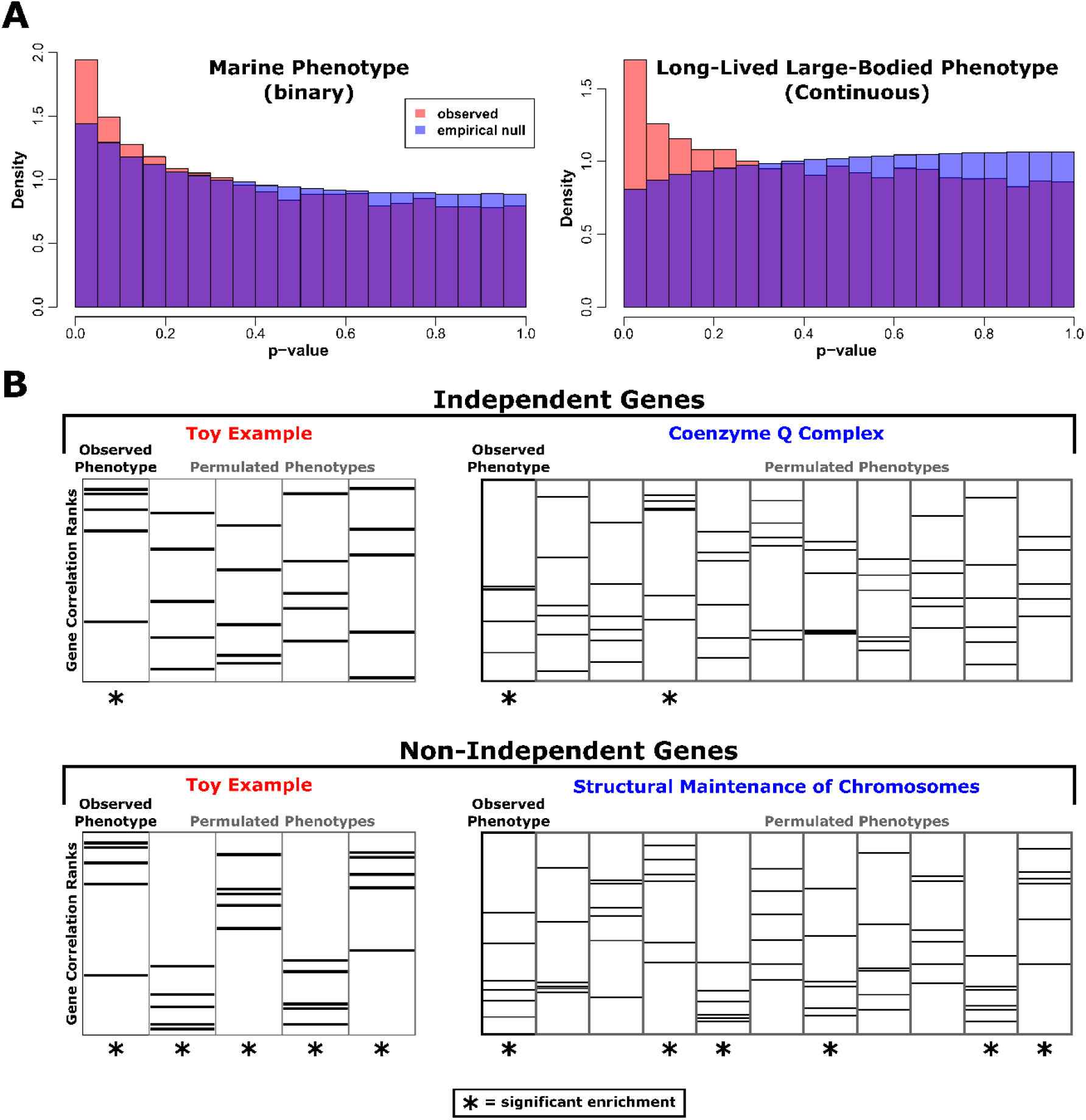
Permulations reveal statistical anomalies in gene- and pathway-level analyses. (**A**) p-value histograms comparing observed/parametric p-values to the empirical null p-value distribution from permulations for a binary phenotype (marine) and a continuous phenotype (long-lived large-bodied). In both cases, the empirical null (shown in blue) is non-uniform. As a result, for the binary phenotype genes have artificially inflated significance and for the continuous phenotype genes have artificially deflated significance. (**B**) Pathway enrichment statistics demonstrate artificially inflated significance because genes in many pathways are non-independent. Permulations correct for non-independence by quantifying the frequency at which significant pathway enrichment occurs due to chance.

#### Problem 2: Non-independence among genes gives rise to false positives in pathway enrichment analyses

Standard pathway enrichment analyses, such as those implemented in GOrilla and GO::TermFinder, interrogate whether whole groups of functionally related genes are over-represented among a statistically determined subset of ranked list (Boyle et al. 2004; Eden et al. 2007; Eden et al. 2009). These methods can be easily applied to the output of evolutionary rate analysis in the hope that they will yield insight into which pathways and functional annotations are enriched among convergently evolving genes. However, in the case of measuring evolutionary rates, it is clear that genes are not independent because functionally related genes experience correlated evolutionary rate shifts both due to interacting protein products and changes in constraint on their shared function (Juan et al. 2008; Clark et al. 2012; Clark et al. 2013). Therefore, many functionally related genes “travel in packs” in association with a phenotype (**Figure 1B**). In other words, if one gene in a group appears to be associated with a phenotype, the other genes in the group will as well because of their connection to the first gene. The result is that a function could appear as associated with the phenotype due to actual involvement, but also potentially by random chance, causing an erroneous inference of enrichment. Because the genes travel in packs, simple enrichment tests assign undue confidence to an essentially spurious enrichment.

To tackle these problems, we have developed a novel strategy that combines *permu*tations and phylogenetic simu*lations* to generate null phenotypes, termed “permulations”. The strategy addresses the issue of a complex null empirically by generating simulation-based permutations that account for the phylogenetic information in the observed phenotype, both for the binary and continuous contexts. It also more accurately mimics the null expectation for a given phenotype by exactly matching the distribution of observed phenotype values, for continuous phenotypes, and exactly matching number and structure of foreground branches, for binary phenotypes. We use these permulated phenotypes to calculate empirical p-values for gene-phenotype associations and pathway enrichment related to a phenotype. In doing so, we have created a statistical pipeline that accurately reports confidence in relationships between genetic elements and phenotypes at the level of both individual elements and pathways.

### New Approaches

#### Permulation: A Hybrid Approach of Using Permutation and Phylogenetic Simulation to Generate Null Statistics

The goal of permulations is to empirically calibrate p-values from phylogenetic methods by producing permutations of the phenotype tree that account for the structure in the data, resulting in what we term “permulation p-values”. The permulation method requires a master species tree and a species phenotype (either continuous or binary). The method then returns a set of phenotypes that are random but preserve the phylogenetic dependence of the input phenotype.

We typically generate 1,000 such permulated phenotypes which are then used in the RERconverge framework to compute gene-trait associations, resulting in 1,000 empirical null statistics for each gene. Similarly, we can also run enrichment analyses using the permulated phenotypes to produce 1,000 empirical null statistics for each pathway. Finally, for each gene or pathway, we calculate the permulation p-value as the proportion of empirical null statistics that are as extreme or more extreme than the observed parametric statistic for that gene or pathway. Since empirical null statistics capture the true null distributions for genes and pathways, the permulation p-values represent the confidence we have to reject the null hypotheses of no association, correlation, or enrichment given the underlying structure of our data. Our permulation methods for binary and continuous phenotypes have been included in the publicly available RERconverge package for R (Kowalczyk et al. 2019) (published on github at https://github.com/nclark-lab/RERconverge), with a supplementary walkthrough (see **Supplementary Walkthrough**) also available as a vignette included in the RERconverge package.

#### Phylogenetic Permulation for Continuous Phenotypes

For continuous traits, the permulation is a two-step process. Given the master tree, representing the average evolutionary rates across all genetic elements, and phenotype values for each species, we simulate a random phenotype using Brownian motion model. The simulated values are then assigned the real phenotype values based on ranks. The species with the highest simulated value is assigned the highest observed value, the species with the second-highest simulated value is assigned the second-highest observed value, and so on. By doing so, observed phenotypes are shuffled among species with respect to the underlying phylogenetic relationships among the species, so more closely related species are more likely to have more similar phenotype values than more distantly related species. (**Figure 2**). In the RERconverge package, the function *simpermvec* generates a permulated phenotype given the original phenotype vector and the underlying phylogeny with appropriate branch lengths. The master tree from the RERconverge *readTrees* function is appropriate to use for simulations. In most cases, the user will not have to use the *simpermvec* function directly — instead, the *getPermsContinuous* function that calculates null empirical p-values for genes correlations and pathway enrichments will call *simpermvec* internally.

**Figure 2.**
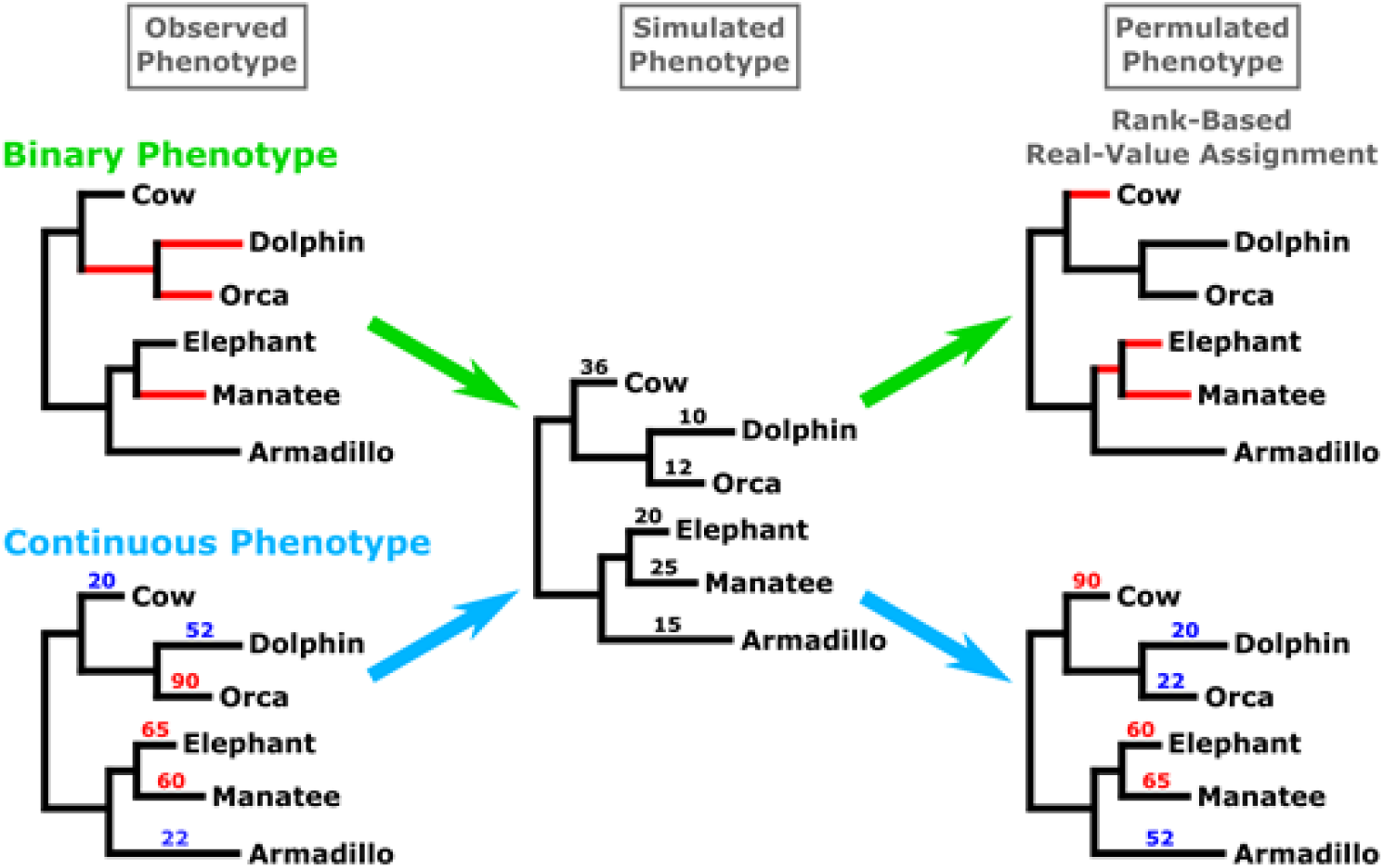
Permulated phenotypes were generated by simulating phenotypes and then assigning observed phenotype values based on the rank of simulated values. Simulations were performed using Brownian motion phylogenetic simulations and a phylogeny containing all mammals with branch lengths representing the average evolutionary rate along that branch genome-wide. For binary phenotypes, foreground branches for permulated phenotypes are assigned based on the highest-ranked simulated values while preserving the phylogenetic relationships between foregrounds. For continuous phenotypes, observed numeric values were assigned directly to species based on ranks of simulated values.

#### Phylogenetic Permulation for Binary Phenotypes

For binary traits, the critical feature is the number of foreground species and their exact phylogenetic relationship, and hence the inferred number of phenotype-positive internal nodes or equivalently phenotypic transition. The two-step process proposed above does not guarantee to perfectly preserve this structure. Instead we employ a rejection sampling strategy where the simulation is used to propose phenotypes which are accepted only if they match the stricter requirements. Using the simulation as the proposed distribution ensures that phylogenetically dependent phenotypes are generated and thus speeds up the construction of null phenotypes over what can be achieved from random selection. Specifically, the simulation outcome is used to choose the same number of binary foreground species and preserve the phylogenetic relationships among chosen foregrounds, as observed in the actual foregrounds (**Figure 2**, Binary Phenotype).

We present two binary permulation strategies that account for requirements of different software packages for the topology of the input phenotype tree. The strategies also encompass the tradeoff between computational feasibility and statistical exactitude—in some cases, it may not be possible to perform one method, in which case the other method is a viable alternative. The complete case (CC) method is the first and simpler strategy. The CC method performs permulations using the master tree in which all species are present and therefore generates permulated trees that contain the complete set of species. Since the CC method produces one set of permulated phenotypes for all the genes, the exact number of foreground and background species per genetic element may not be preserved because of species presence/absence in those alignments (**Figure 3**). Thus, the CC method is an imperfect but fast method to generate null phenotypes, but we recommend use of the SSM method whenever feasible.

**Figure 3.**
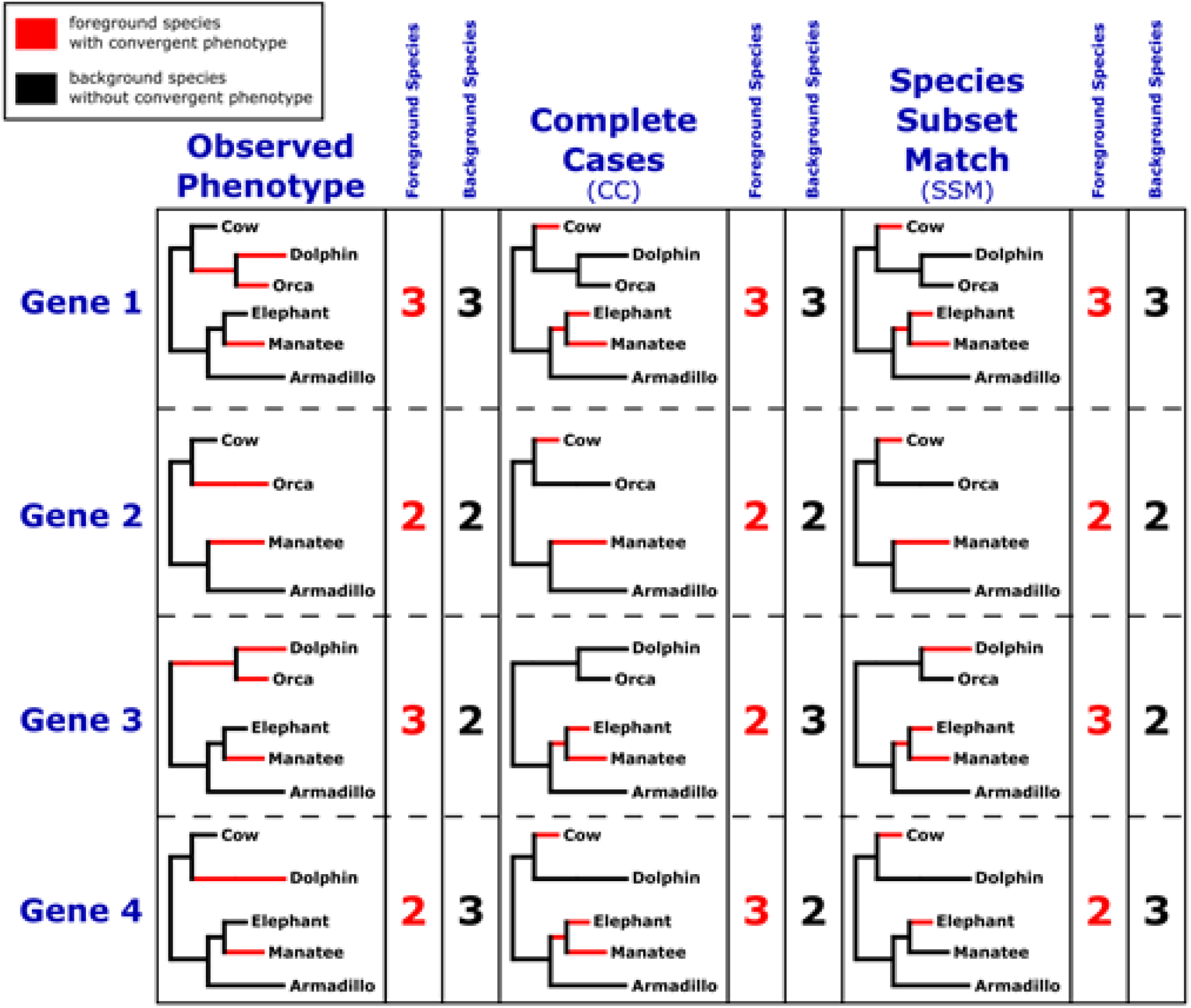
When permulating binary phenotypes, missing species are handled using either the complete case (CC) method or the species subset match (SSM) method. For the CC method, top-ranked simulated values are assigned as foreground regardless of gene-specific species absence. For the SSM method, top-ranked simulated values are assigned as foreground after considering gene-specific species absence so the number of foreground and background species for each gene is consistent across every permulated phenotype. Note that in the case of genes with all species present (e.g., Gene 1), CC and SSM methods are identical.

In contrast, the species subset match (SSM) method accounts for the presence/absence of species in different gene trees. For each permulation, the SSM method generates separate null phenotypes for each tree in the set of genetic elements. Since genetic element-specific trees contain exactly the species that have that genetic element, the null phenotypes exactly match the observed phenotypes for that genetic element in terms of number of foreground and background species (**Figure 3**). Additionally, unlike the CC method, null phenotypes for a single permulation iteration are distinct, and potentially unique, from each other because they are generated on a genetic element-by-genetic element basis. Although the SSM method is statistically more ideal than the CC method, it is much more computationally intense and may not be feasible for very large datasets. In RERconverge, CC and SSM permulations can be performed using the *getPermsBinary* function, setting the “permmode” argument as “cc” and “ssm”, respectively.

## Results

### Permulation of Binary Phenotypes Improved Power and Type I Error Control

To evaluate the performance of the permulation methods compared to the parametric method for binary phenotypes, we use RERconverge to find genetic elements that demonstrate convergent acceleration of evolutionary rates in response to the marine environment. For all the ensuing analyses, we use the set of protein-coding gene trees across 63 mammalian species previously computed in (Partha et al. 2019). These trees have the “Meredith+” tree topology (Kowalczyk et al. 2020) (**Figure 4**), a modification of the tree topologies published in (Meredith et al. 2011) and (Bininda-Emonds et al. 2007), resolved for their differences across various studies as originally reported in (Meyer et al. 2018). We set marine mammals evolving from three independent lineages as foregrounds (blue branches in **Figure 4**): pinnipeds (Weddell seal, walrus), cetaceans (bottlenose dolphin, killer whale, the cetacean ancestor), and sirenians (West Indian manatee) (Chikina et al. 2016). We consider three p-value calculation methods: parametric, complete case (CC) permulations, and species subset match (SSM) permulations. Resulting p-values are corrected for multiple hypothesis testing using Storey’s correction (Storey and Tibshirani 2003; Storey et al. 2020).

**Figure 4.**
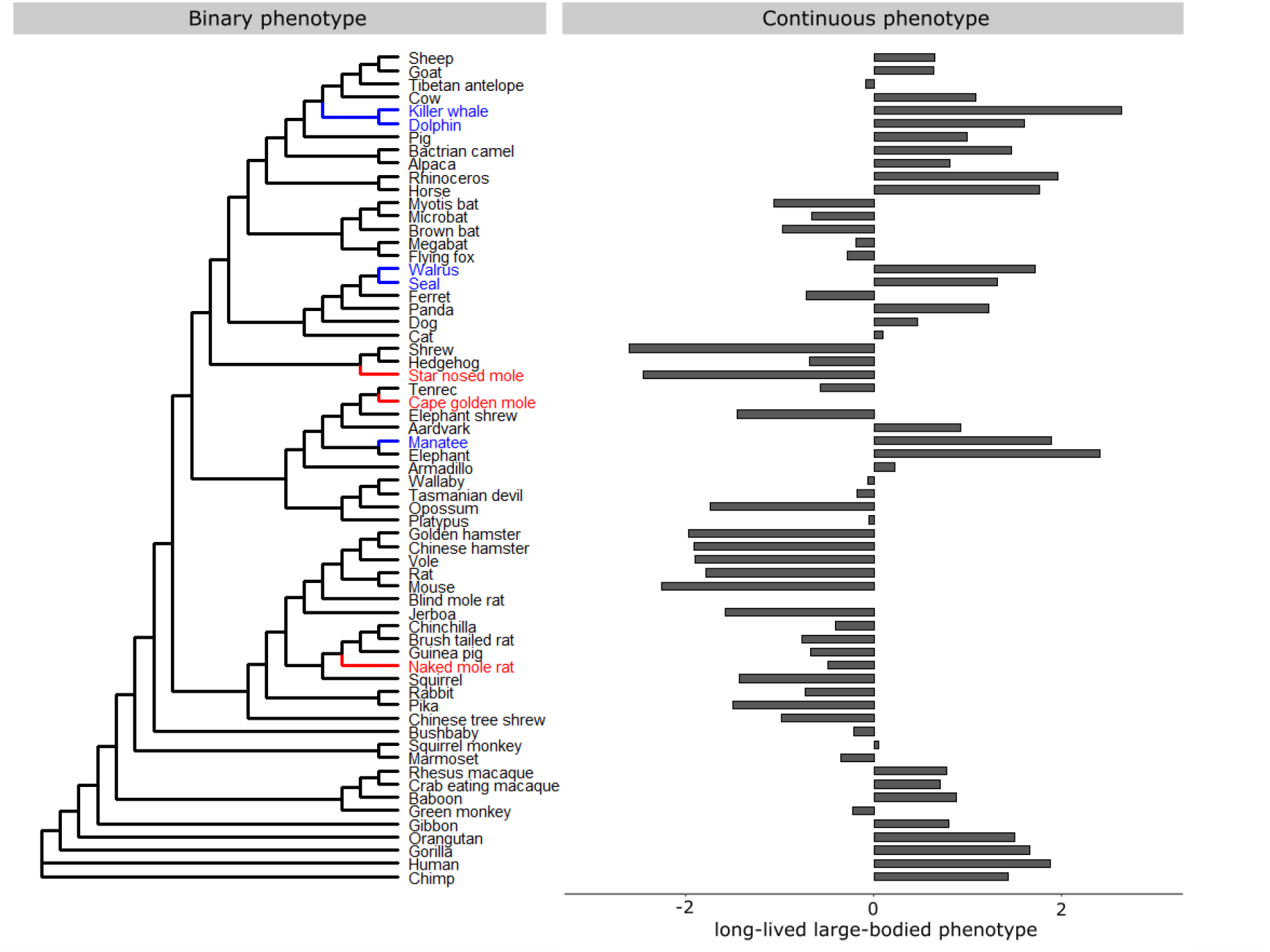
Meredith+ tree topology and the binary and continuous phenotypes evaluated. Binary phenotypes include the marine mammal phenotype and the subterranean mammal phenotype (foreground branches are indicated in blue and red, respectively). The continuous phenotype evaluated is the long-lived large-bodied phenotype as constructed based on the first principal component between species body size and maximum longevity (Kowalczyk et al. 2020).

We see in **Figure 1A** that the parametric p-values for the association of genes with the observed marine phenotype (red histogram) are enriched with small p-values. According to the standard parametric approach, which assumes a simple null hypothesis with uniformly distributed p-values, the enrichment of low p-values indicates the possible presence of genes with evolutionary rate shifts that are significantly correlated with marine adaptation. However, when we construct the empirical null p-value distribution by permulating the marine phenotype, we see that the null distribution is not uniform. In fact, the enrichment of low p-values is also present in the null distribution (blue histogram), albeit a lesser enrichment than the observed, meaning that observing low p-values by chance is more likely than expected. Thus, if we use standard multiple testing procedures directly on the parametric p-values, we will identify more positive genes than the true number of positives, in other words causing an undercorrection of p-values.

In order to demonstrate that our permulation method effectively corrects for the background p-value distribution, we plot histograms of parametric and permulation p-values obtained from the parametric and permulation methods, respectively (**Figure 5A**). Compared to the parametric method, the histograms for the CC and SSM permulations have steeper slopes at low p-values, indicating that the permulation methods have better Type I error control. Furthermore, the histograms for the permulation methods plateau at higher π_0_ than the parametric method, consistent with the postulation that the parametric method would reject more (possibly false) positives. These findings are also observed when we identify genes with significant evolutionary acceleration in marine mammals by setting a rejection threshold of Storey’s FDR ≤ 0.4 (the high threshold is set considering the high minimum FDR from the parametric method), as is shown in **Figure 5B**. For the permulation methods, as the number of permulations increases, the number of identified marine-accelerated genes increases and eventually stabilizes after ~400 permulations. The asymptotic numbers of marine-accelerated genes identified by permulations (~350 genes for CC permulation and ~450 genes for SSM permulation) are much smaller than the ~700 genes identified through parametric statistics, demonstrating improved Type I error control.

**Figure 5.**
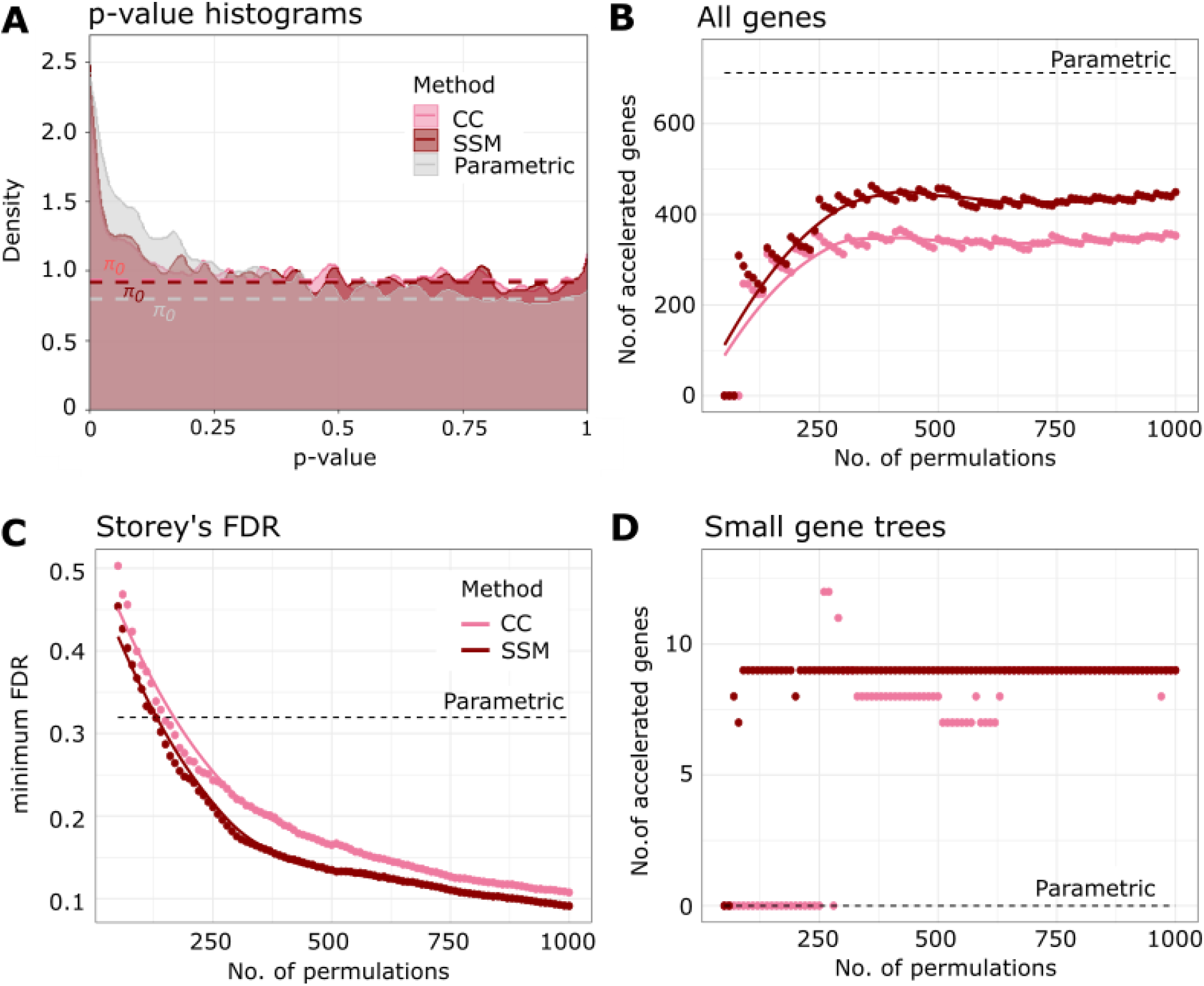
Permulation of binary phenotypes corrects for inflation of statistical significance in finding evolutionarily accelerated genes in marine mammals. (**A**) Histogram of parametric and permulation p-values from the parametric, complete case (CC) permulation, and species subset match (SSM) permulation methods. (**B**) Permulation methods identify fewer accelerated genes in marine mammals compared to the parametric method, correcting for the inflation of significance. The rejection region of the multiple hypothesis testing is set to be Storey’s FDR ≤ 0.4, considering the weak power of the parametric method. (**C**) Binary permulation methods have greater statistical power compared to the parametric method, as shown by the minimum false discovery rate (FDR) calculated using Storey’s method. (**D**) Permulation methods can identify accelerated genes that are missing in many species (gene tree size ≤ 30), whereas the parametric method fails to do so.

Surprisingly, while the permulation methods reject fewer regions, we have greater confidence in these rejections. **Figure 5C** shows the minimum corrected p-values achieved by the permulation methods with increasing number of permulations. The figure shows that the permulation methods provide better control of FDRs compared to the parametric method with only a few permulations (above ~125 permulations). With increasing permulations, the minimum FDRs continues to drop to reach levels below 0.1 at 1000 permulations, while the minimum FDR from parametric statistics is higher at above 0.3 for Storey’s correction. Use of the permulation null significantly improves the statistical power of the method and provides much higher confidence in detecting true correlations between evolutionary rate shifts and the convergent phenotype of interest.

Lastly, we find that permulation methods can identify marine-accelerated genes that are missing in many species, i.e., genes with phylogenetic trees containing few species. In contrast, the parametric method fails to reject any such gene (**Figure 5D**).

### Binary Permulation Methods Improved Gene-level Detection of Functional Enrichment

We have demonstrated that the permulation methods show favorable statistical properties based on the distribution of p-values. We expect that this approach also improves the biological signal of rate convergence analysis. In order to address this question, we ask if the marine-accelerated genes identified by binary permulations are enriched with functions that are consistent with the marine phenotype. Our group previously identified marine-specific pseudogenes that should be undergoing accelerated evolution in marine mammals due to relaxation of evolutionary constraint (Meyer et al. 2018). Putative pseudogenes associated with marine mammals were identified using Bayes Traits software (Pagel and Meade 2006) to find signals of coevolution between marine status and pseudogenization. In addition, our group also previously found that many marine-accelerated genes that evolve under relaxed constraint are enriched with genes responsible for the loss of olfactory and gustatory functions (Chikina et al. 2016). Thus, to represent the “ground truth”, we select a collection of gene sets relevant to olfactory and gustatory functions from the Mouse Genome Informatics (MGI) database and top-ranking marine-specific pseudogenes with Bayes Traits FDR values less than 0.25.

We then perform the one-tailed Fisher’s exact test to measure the enrichment of the functions in the marine-accelerated genes from the parametric and permulation methods. The Fisher’s exact test odds ratios indeed show that the CC and SSM permulation methods generally magnify or maintain the effect sizes of enrichment across the gene sets compared to the parametric method (**Figure 6A**). At worst, the permulation methods match the performance of the parametric method (e.g., “taste/olfaction phenotype” gene set). The improved performance of the permulation methods is also demonstrated in the example precision-recall curves for the marine-associated pseudogenes in **Figure 6B**.

**Figure 6.**
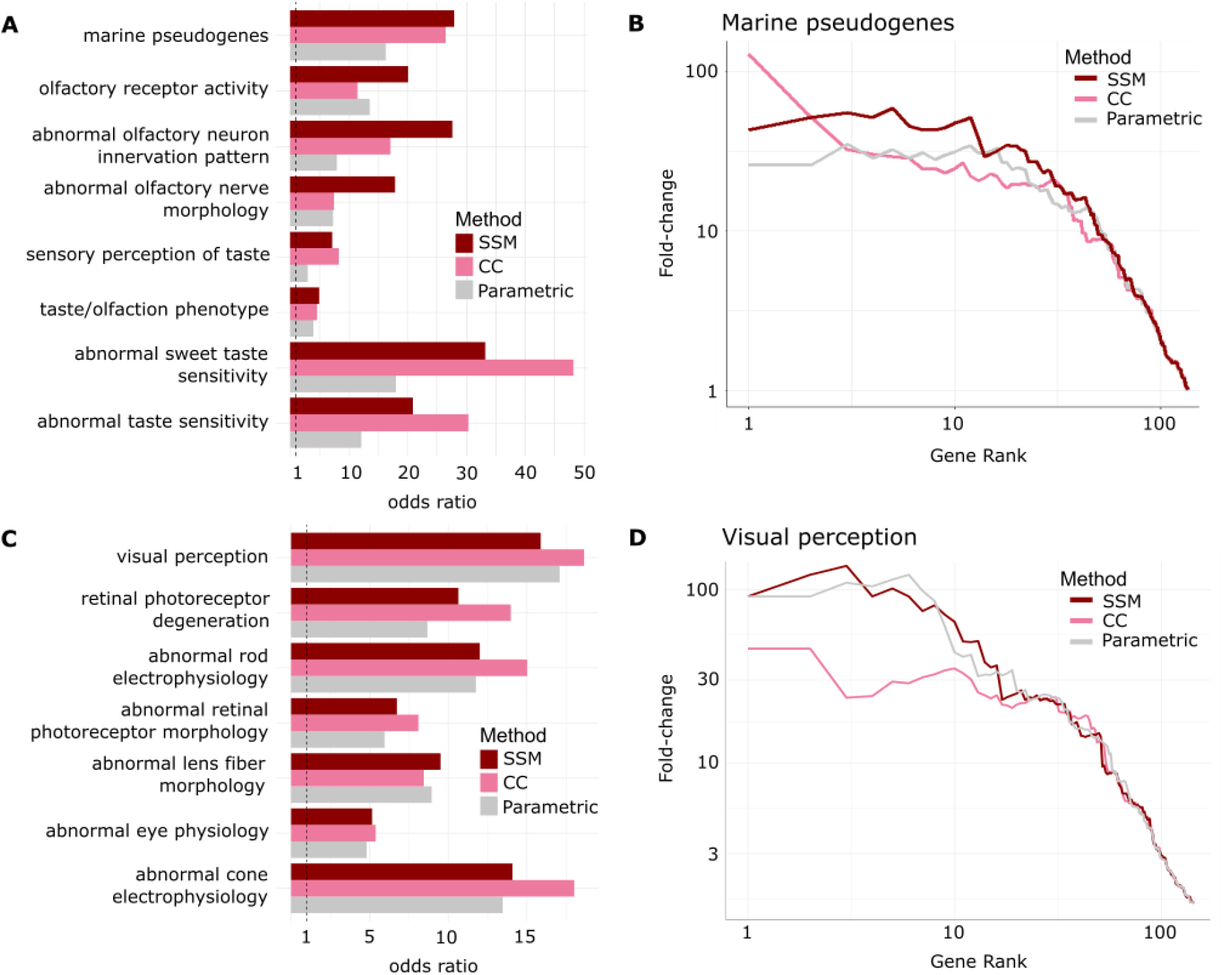
Binary permulation methods have matching or improved power compared to the parametric method in detecting enrichments of functions consistent with known phenotypes. (**A**) Fisher’s exact test odds ratios showing that marine-accelerated genes identified by the permulation methods have greater enrichment of gustatory and olfactory genes, compared to the parametric method. (**B**) Precision-recall curves for the enrichment of the marine pseudogenes in the identified marine-accelerated genes. (**C**) Fisher’s exact test odds ratios showing that subterranean-accelerated genes identified by the permulation methods have greater or comparable enrichment of ocular genes, compared to the parametric method. (**D**) Precision-recall curves for the enrichment of the visual perception genes in the identified subterranean-accelerated genes.

To see if this observation generalized to other phenotypes, we repeat the whole analysis to find genes that are accelerated due to subterranean adaptation, defining three independent mole species (naked mole rats, star-nosed moles, and cape golden moles), for which high quality genomes are available in our dataset, as foreground species (red branches in **Figure 4**). As subterranean-accelerated genes have been found to be enriched in ocular functions (Prudent et al. 2016; Partha et al. 2017; Partha et al. 2019), we pick gene sets relevant to vision-related functions as the “ground truth”. In general, the signals we obtained from RERconverge for this phenotype are much weaker than in the marine phenotype case, but the enrichment is still captured in the rankings of the genes. Similar to the marine phenotype, permulation methods generally improve or match the performance of the parametric method (**Figure 6C** and **D**).

### Binary Permulation Method Corrects for False Positives in a Related Approach

We apply CC permulations to Forward Genomics, an alternate method that tests for an accelerated evolutionary rate in a set of foreground species. The SSM method is not tested because Forward Genomics does not allow for unique foreground specification across individual genes, but instead uses one set of foreground species across all genes. We generate 500 CC-permulated trees, but because Forward Genomics only works for tree topologies where there are at least 2 foreground species, the true number of permulations tested was 417, as some of the permulated trees did not contain 2 or more trait loss species.

Forward Genomics’ “global method” uses substitution rate with respect to each tree’s root to correlate with trait loss and identify convergent relaxed selection; therefore, it does not correct for evolutionary relatedness. The “local branch method”, an improvement on the original approach, uses substitution rate with respect to the most recent ancestor to identify relaxed selection, which substantially improves its power (Prudent et al. 2016). We used both methods to demonstrate that applying a permulation correction to each method’s p-values improves performance even for a statistic with known flaws.

Both the global and local methods had unusual p-value distributions. The local method identified high proportion of positives with significant p-values (**Figure 7A**), while p-values from the global method were highly concentrated around 0.5 (global p-values not shown). Adjusting for multiple testing further exaggerated this issue. For the global method, due to the number of genes with very low p-values, the lowest possible Benjamini-Hochberg (BH) corrected parametric p-value is 0.531, and for the local method, the lowest possible corrected p-value is 0.4647. For the local method, out of 18,797 genes, more than half of the genes (12,438) have the lowest possible corrected parametric p-value. As such, it is impossible to designate a significance cut-off, because it will either include no genes or include most of the genes. Applying the permulation strategy to Forward Genomics, output we find that of the same set, 889 have a corrected permulation p-value that is less than or equal to 0.465 (the minimum observed), allowing for a more reasonable selection of a rejection threshold.

**Figure 7.**
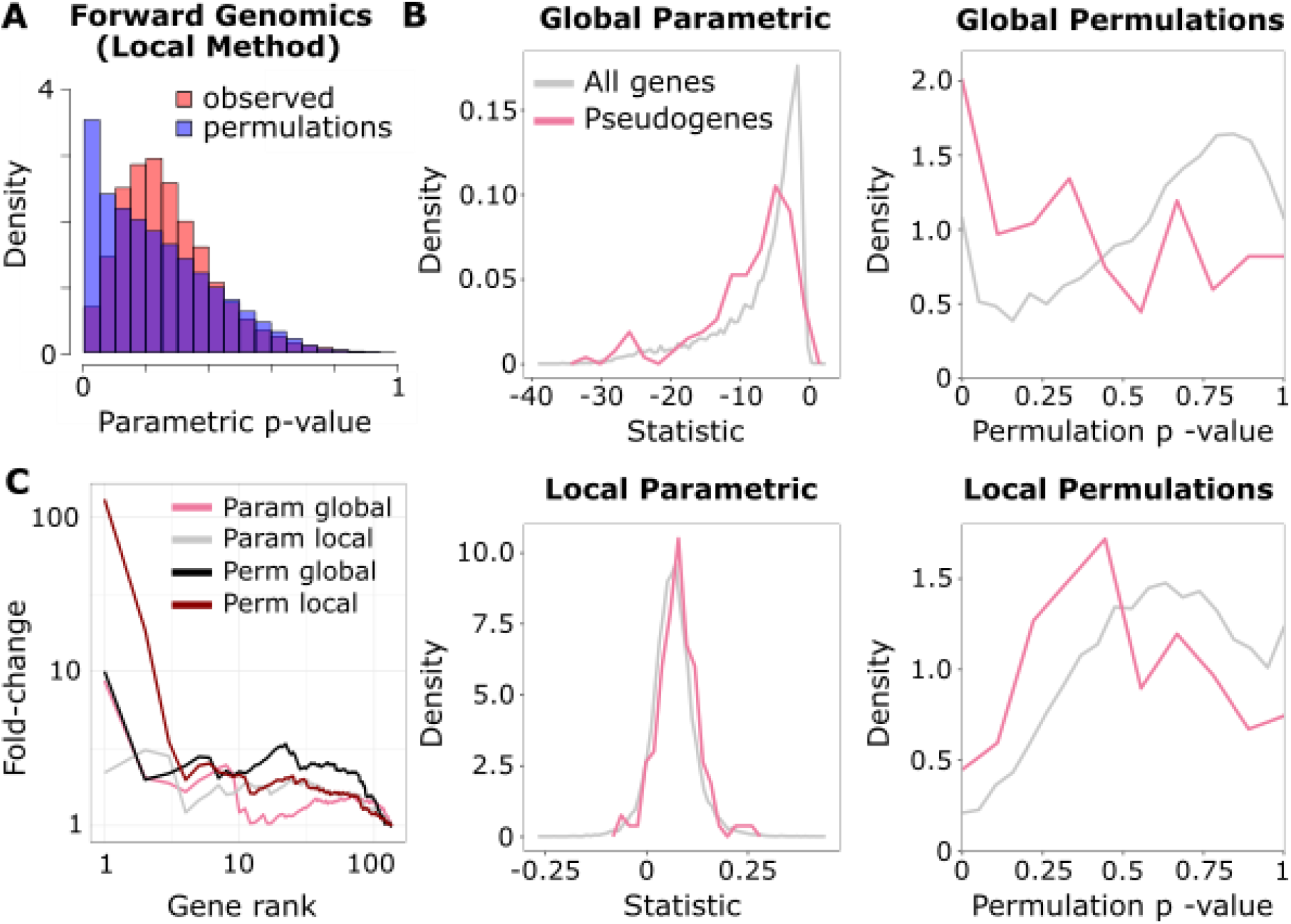
Binary permulation methods improve Forward Genomics’ positive-predictive value and power. (**A**) Distribution of the empirical null p-value and the parametric p-value for Forward Genomics’ local method. Note that the empirical null distribution (in blue) is highly non-uniform and observed parametric p-values are strongly right skewed, both of which increase false positive rate. (**B**) Distributions of Forward Genomics statistics and corresponding permulation p-values for local and global methods. Both global and local statistics show slight shifts (to the left for global statistics and to the right for local statistics) indicating enrichment of marine mammal pseudogenes under accelerated evolution (global AUC=0.6235; local AUC=0.6196). Permulation p-values show a more dramatic shift toward significant values for marine pseudogenes under accelerated evolution for the global method (AUC=0.6653) and about the same shift for the local method (AUC=0.6086) compared to parametric statistics. (**C**) Precision-recall curves for the enrichment of pseudogenes in marine-accelerated genes using parametric statistics and permulation p-values for both local and global methods. Permulated values represent a unique ranking in which ties in permulation p-values for genes are broken based on parametric statistics.

In addition to analyzing results at the individual gene level, we also used the marine pseudogenes as a “ground truth” set of genes that should be undergoing accelerated evolution in marine species, to test our ability to detect these genes. As shown in **Figure 7B**, the global and local parametric test statistics show slight enrichment for elements that are pseudogenized in marine mammals, and the difference is improved when permulation p-values are computed. **Figure 7C** shows the same data as a precision-recall plot, clearly demonstrating that the permulation correction improves the predictive power of both methods.

### Permulations Improve Power to Detect Genes Correlated with a Continuous Phenotype

To evaluate the permulation strategy for continuous phenotypes, the long-lived and large-bodied phenotype is used as defined in previous work (Kowalczyk et al. 2020). The numerical phenotype is constructed by calculating the first principal component between body size and maximum lifespan across 61 mammal species (**Figure 4**, Continuous Phenotype). The first principal component therefore represents the agreement between body size and lifespan – species like whales with long lifespans and large sizes have large phenotype values and species like rodents with short lifespans and small sizes have small phenotype values.

One thousand permulations are performed to generate 1,000 null statistics and p-values for each gene, as well as to calculate permulation p-values as the proportion of null statistics that were as extreme or more extreme than the observed statistic per gene. As shown in **Figure 1A**, the empirical null p-value distribution for genes associated with the long-lived large-bodied phenotype is non-uniform, and in fact slopes down at low p-values. This indicates that observing small p-values due to chance alone happens less often in our dataset than we would typically expect compared to the standard uniform expectation. In practice, the result of the non-uniform null is overcorrection of parametric p-values using a standard multiple hypothesis testing correction. In other words, for this dataset, corrected parametric p-values are larger than they should be when using multiple hypothesis testing correction (such as a Benjamini-Hochberg correction) that assumes a uniform null. The null distribution of permulation p-values, however, does follow a standard uniform null, so Benjamini-Hochberg corrected permulation p-values represent our true, higher confidence in a correlation between gene evolutionary rate and phenotypic evolution. We observe this increased confidence in our data – after multiple hypothesis testing correction, only 24 parametric p-values remain significant at an alpha threshold of 0.15 while 305 permulation p-values remain significant. Regardless of the increase in power, permulation p-values provide a more accurate representation of confidence in rejecting the null hypothesis, and thus are a more valid metric than parametric p-values.

### Permulations Correct Pathway Enrichments for Genes with Correlated Evolutionary Rates

After generating null p-values and statistics from permulations for either binary or continuous traits, those values can be used to calculate null pathway enrichment statistics. Permulation p-values for pathways are then calculated as the proportion of null pathway enrichment statistics as extreme or more extreme than the observed statistic. This procedure corrects for gene sets with correlated evolutionary rates, that is genes whose rates will “travel in packs” regardless of any relation to the phenotype (**Figure 1B**). Such groups of genes will tend to show enrichment more often than would be observed if the genes rates were independent after conditioning on phenotype, resulting in false signals of pathway enrichment.

**Figure 1B** demonstrates how correlated evolutionary rates can cause problems in pathway enrichment analyses. Each vertical bar represents all genes in the genome and horizontal black lines represent genes in a pathway of interest. Genes are ranked based on gene-phenotype associations, so clusters of genes either at the top or bottom of the lists indicates enrichment. Different vertical bars represent either the observed phenotype or different permulated phenotypes. In the case of independently evolving genes, the typical null expectation holds—permulated phenotypes show a random distribution of ranks with non-significant enrichment statistics. On the other hand, in the case of genes that do not evolve independently, the typical null expectation does not hold. Even when using a fabricated phenotype (a permulation phenotype), genes appear to cluster at the extremes of the ranked list. The clustering, and resulting enrichment, is caused by the genes “traveling in packs” – if one gene in a pack is associated with a phenotype, all the genes in that pack will appear to be associated with the phenotype because they are not truly independent observations.

This phenomenon is well described in the context of gene expression and is typically handled by performing label permutations (Subramanian et al. 2005; Majewski et al. 2010; Ritchie et al. 2015) and in certain cases parametric adjustments (Wu and Smyth 2012). However, simple label permutations are not applicable to associations involving a phylogeny as they would not preserve the underlying phylogenetic relationships, thereby producing false positives. Our permulation strategy avoids this pitfall by sampling permutations from the correct covariance structure that captures the underlying phylogenetic dependence.

Permulations account for the non-independence problem by explicitly incorporating it into the null distribution used to calculate permulation p-values. In the demonstrated case of the Coenzyme Q Complex, only one permulation out of the ten depicted shows enrichment due to random chance (indicated by an asterisk * below the vertical bar in **Figure 1B**), which would correspond to a permulation p-value of 0.1 in this toy example. This interpretation is identical to the standard p-value interpretation—the proportion of times we expect to see a statistic as extreme or more extreme than observed *assuming that the null expectation is true*. In the case of permulations, we simply explicitly calculate the null expectation rather than using a predefined distribution (t-distribution, F-distribution, etc.). In the case of enrichment for a pathway with non-independent genes, the significance of the permulation p-value will agree with the significance of the parametric p-value because the null expectation from permulations agrees with the typical null expectation.

In the case of a pathway with genes with non-independent evolutionary rates, the permulation p-value will be larger than the parametric p-value because the permulation p-value will penalize for non-independence. An example with “Structural Maintenance of Chromosomes” genes shows that, although there is an apparent enrichment based on the observed phenotype, half (5 out of 10) of permulated phenotypes show at least as strong enrichment for a permulation p-value of 0.5. Therefore, although the pathway does appear to be enriched from parametric statistics, its enrichment is actually not exceptional given the null expectation for that set of genes.

Permulation p-values are calculated for every pathway individually. **Table 1** shows top enriched pathways under accelerated evolution and decelerated evolution in association with the long-lived large-bodied phenotype. While most significantly enriched pathways under decelerated evolution based on parametric p-values also demonstrate significant permulation p-values, many pathways under significant acceleration show non-significant permulation p-values. Thus, this phenotype shows little evidence for accelerated pathway evolution associated with phenotypic evolution.

**Table 1.**
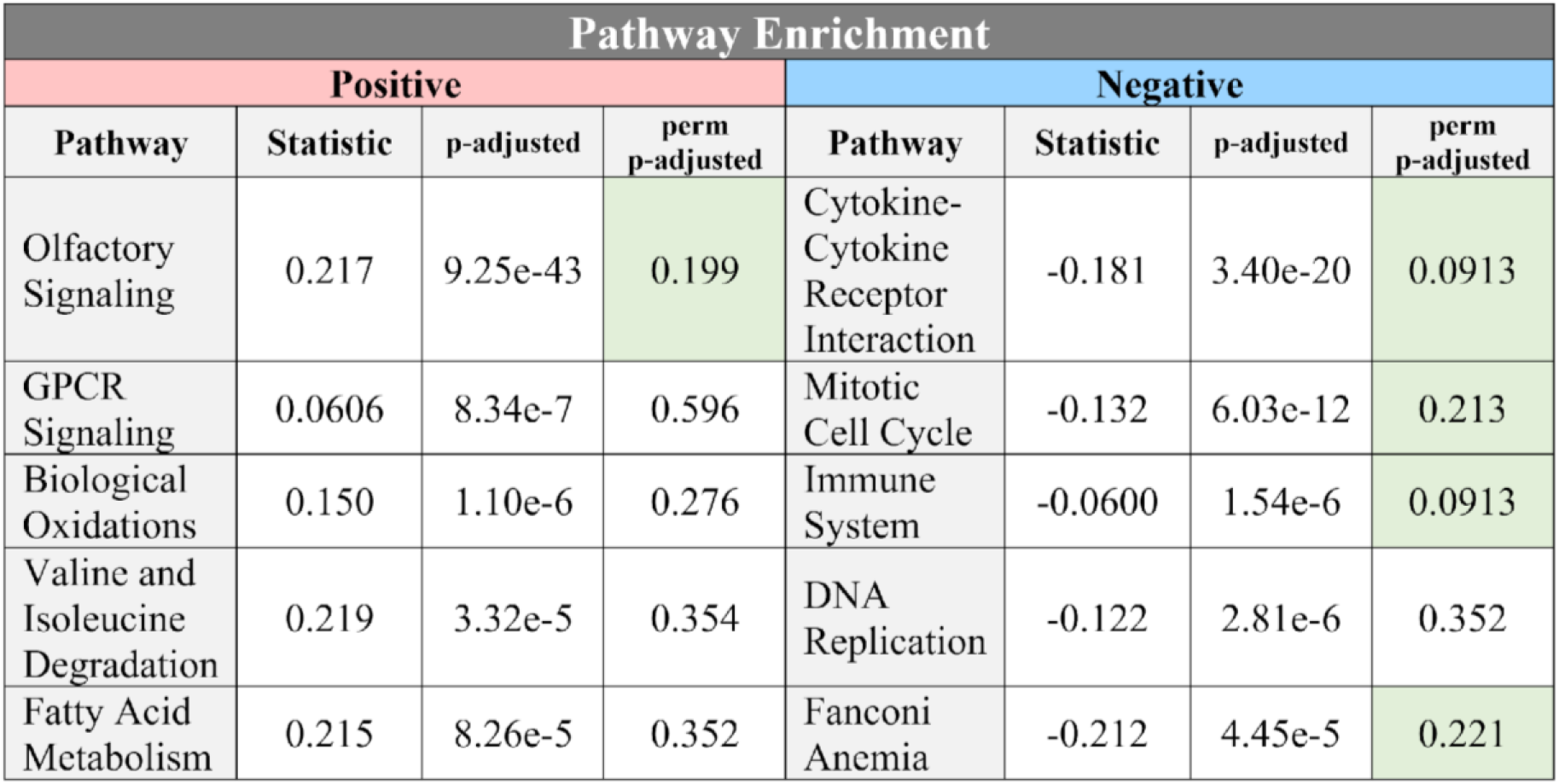
Top-enriched pathways with quickly evolving genes and slowly evolving genes in association with the long-lived large-bodied phenotype according to parametric p-values. Note that due to the number of pathways, the lowest possible Benjamini-Hochberg corrected permulation p-value is 0.0913. Boxes in green show significance at alpha = 0.25. Note that many accelerated pathways that appear to be enriched based on parametric p-values are not enriched based on permulation p-values.

### Comparison of Phylogenetic Simulations, Permutations, and Permulations

At the pathway level, permulations result in p-values that are about equally as conservative as phylogenetic simulations alone and more conservative than permutations alone (**Figure 8**). Both permulations and simulations are preferred to permutations because null phenotypes generated from permulations or simulations reflect the underlying phylogenetic relationships among species, while null phenotypes from permutations do not. Therefore, the empirical null generated from permulations or simulations more closely represents the true null expectation for phenotype evolution. Although permulations and simulations show similar performance, we prefer permulations because permulated phenotypes exactly match the distribution of observed phenotypes, and thus create null phenotypes uniquely tailored to a particular continuous phenotype of interest. Such matching eliminates statistical anomalies that can arise due to discrepancies in range and distribution of permulated phenotypes compared to observed phenotypes.

**Figure 8.**
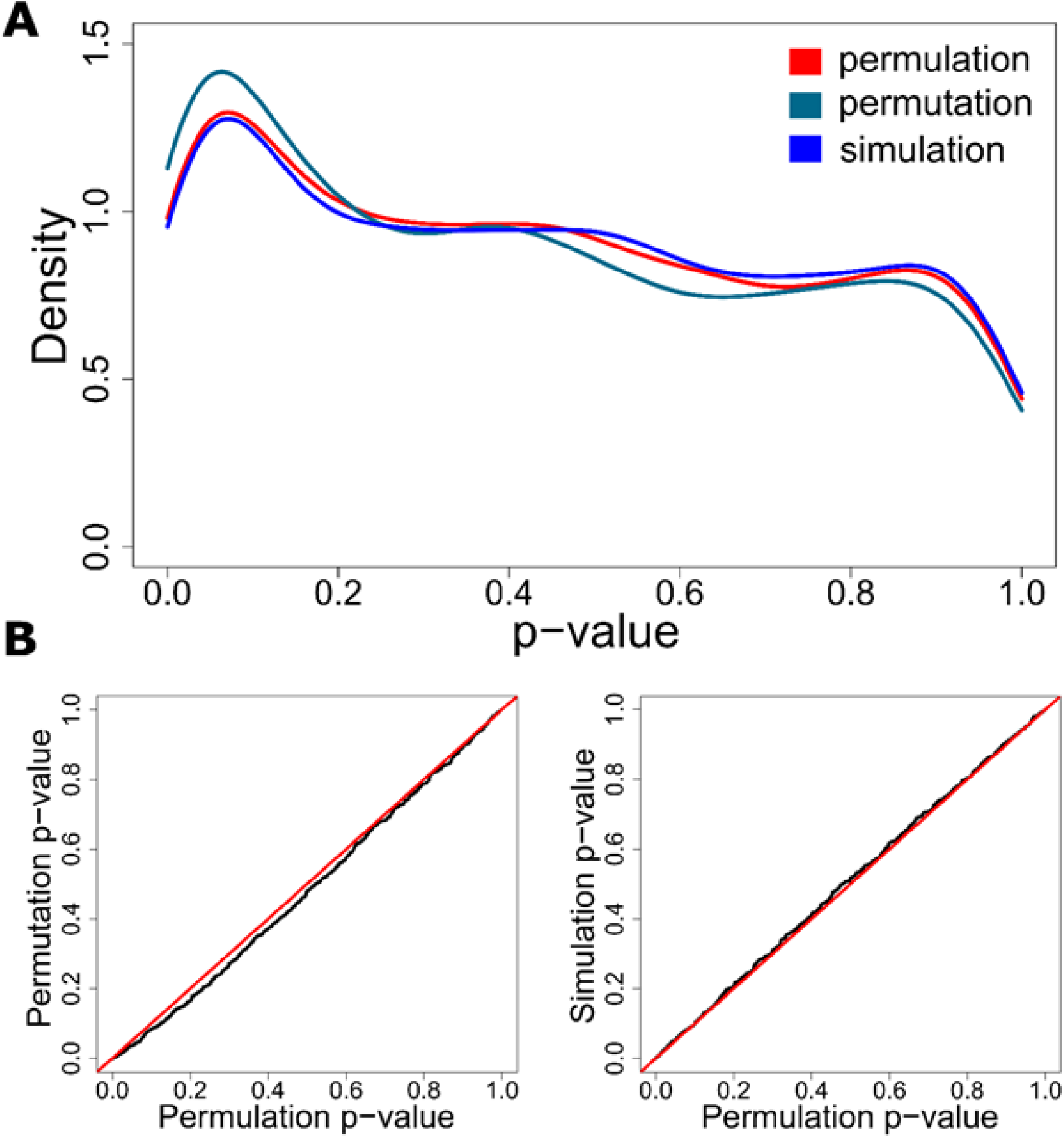
Permulations p-values are more conservative than permutation p-values and about equally as conservative as simulation p-values. All plots demonstrate enrichment for canonical pathways associated with the long-lived large-bodied phenotype. (**A**) Density plots representing the empirical p-value distributions for the three methods to generate null p-values. Permulation and simulation curves are very similar, while the permutation curve demonstrates a stronger enrichment of low p-values and therefore less conservative p-values. (**B**) Q-Q plots comparing empirical p-values from permulations to empirical p-values from simulations and permutations also demonstrate that permulation p-values are more conservative than permutation p-values and about equally as conservative as simulation p-values.

## Discussion

In the present work, we have developed novel empirical methods for addressing atypical statistical behavior in phylogenetic analysis, which we term permulations. The methods use phylogenetic relationships among species alongside known values of an observed phenotype to inform Brownian motion simulations, based on which permuted phenotypes are then generated. By doing so, the methods empirically construct the null distribution, which is possibly composite, and account for this complexity in multiple hypothesis testing. For permulation of binary phenotypes, the phylogenetic characteristics preserved are the number of foreground branches and the underlying relationships among foreground branches. For continuous phenotypes, the exact distribution of phenotype values is preserved in addition to the underlying phylogenetic relationships among species.

From testing the strategy on binary and continuous phenotypes, we find that our permulation strategy is an effective approach for overcoming challenges in multiple testing with composite nulls in comparative phylogenetic studies. We discuss with examples how our binary and continuous permulation methods fix issues of both undercorrection and overcorrection of p-values for specified phenotypes, and subsequently improve the quality and confidence of prediction. Note that although our examples demonstrate the usefulness of permulations, they are not necessarily representative of how empirical null distributions will deviate from the typical null for all phenotypes over all phylogenies for all sets of genetic elements. In fact, we expect permulations to behave differently as those variables change, and thus the best way to determine how permulations will affect a particular data set is to run the permulation analyses.

Devising a systematic solution for such problems is difficult because the causes of complex null distributions in phylogenetic studies are mostly unclear. Non-uniform null distributions could arise from a faulty statistical test, or when there are groups of the p-value that are correlated (Allison et al. 2002; Hu et al. 2010). In our case, one possible source of gene evolutionary rate correlations is when non-independent clusters of interacting proteins coevolve, where a substitution in one protein alters the selective pressure on other proteins it interacts with, causing them to evolve in packs (Fraser et al. 2004). Systematic solutions such as using mixture models have been developed to handle non-independence issues in genomics and phylogenetics studies (Allison et al. 2002; Stone et al. 2011). In such models, the null distribution is explicitly constructed by assuming that the distribution results from a mixture of multiple components with distinct distributions. However, these models require assumptions that each component follows a specific distribution, which may not be accurate. For example, (Allison et al. 2002) assumes that the null distribution is a mixture of multiple beta distributed components. With the lack of understanding of other factors that can play a role in causing the complex null distributions we see in our data (e.g., number of foregrounds versus backgrounds, missing species, highly conserved genes biasing the calculation of master branch lengths, etc.), empirically correcting p-values using permulation methods allows us to circumvent the need to artificially deconstruct this unknown correlation structure in the data.

For binary phenotypes, our permulation methods choose permuted foreground sets by matching the number of foregrounds and their underlying relationships to those observed in the actual phenotype. This approach of defining null phenotypes can be justified by phylogenetic non-independence, a notion that arises from the implications of shared ancestry (Felsenstein 1985). At the time of divergence, closely related species diverging from a common ancestor are likely to experience similar selective pressures as the ancestor, as well as similar genetic predispositions to respond to the selection pressures.

With progressing evolutionary time, the daughter species will evolve independently in response to their respective environments. Such similarities in environmental pressures and genetic predispositions diminish with increasing evolutionary distance between species, meaning that the variance in phenotype values will increase with increasing divergence in evolutionary time. Considering this phylogenetic non-independence and that adaptations to selection pressures are often assumed to be reflected in evolutionary rates, it is reasonable to preserve the pattern of divergence between foreground species to construct hypothetical null phenotypes, in finding correlations between evolutionary rates and phenotypes. It is impossible to pick a new set of foreground branches with perfectly matching divergence times, but matching divergence patterns can serve as a justifiable workaround because the general implications of shared ancestry on phylogenetic non-independence among the new set of foregrounds would apply in a similar way.

We developed two versions of permulation methods for binary phenotypes. The complete case (CC) algorithm produces one permuted phenotype from the master tree to apply for all genes simultaneously, while the species subset match (SSM) algorithm produces distinct permuted trees for each gene, accounting for the differences in species membership in different gene trees. This makes the CC method statistically imperfect. For example, a gene that is missing in some species will have a phylogenetic tree that is missing some branches. As a consequence of producing permuted trees from the master tree that contains all species, the CC method may not conserve the number and relationships of foregrounds across the permulations of the example gene (e.g., genes 3 and 4 in **Figure 3**). In contrast, the SSM method accounts for differences in numbers and patterns of foregrounds among different genes and addresses each gene independently. This means that the SSM method is the ideal implementation of our concept of binary permulations. However, the CC and SSM methods were developed to account for the fact that existing comparative genomics methods take in phenotype inputs in different forms. For example, Forward Genomics requires one phenotype tree to apply for all genes, while HyPhy RELAX requires multiple phenotype trees with matching topology to each gene. Regardless of the statistical flaw, our results demonstrate that applying the CC method on Forward Genomics is beneficial for improving prediction (**Figure 7**). In addition, the CC method is significantly faster than the SSM method because it only produces one permuted tree for each permulation, instead of a heterogeneous set of permuted trees applying to different genes. Therefore, in the case of limited computational resources or very large datasets in which using the SSM method is infeasible, the CC method can serve as a good alternative.

Our results also demonstrate that binary permulations improve the sensitivity of RERconverge to identify significantly accelerated genes that are missing in many species (**Figure 5D**), i.e. genes with small trees. By construct, genes with small trees suffer from lower statistical power compared to genes with large trees (for example, the number of ways to permute a small tree is much fewer compared to a large tree). As such, pooling all the p-values together to perform multiple testing correction unfairly penalizes genes with small trees. Calculating permulation p-values from multiple empirical permutations is a way to correct for this imbalance in power by indirectly incorporating important covariates, which accounts for the number of foregrounds, backgrounds, the ratio and phylogenetic relationship between them. Indeed, the pooled null permulation p-values have a uniform distribution, establishing the validity of applying standard multiple testing methods to identify significant divergence in evolutionary rates. Future work can evaluate if such benefits are similarly observed when applied to other comparative genomics methods.

Permulations grant increased power to detect genes associated with a continuous phenotype as suggested by the shape of the empirical null distribution (**Figure 1**). Since permulation p-values are more conservative than p-values from permutations alone and equally as conservative as p-values from simulations alone, they offer a valid alternative to phylogenetic simulations that exactly preserve the distribution of phenotypes. Importantly, in doing so permulations preserve the exact range of phenotype values, a critical characteristic related to the power of the correlation calculated between gene evolution and phenotype evolution. Thus, permulations more accurately match the power between observed and permulated statistics compared to observed and simulated statistics.

Although many of our tests of the permulation strategy were performed using RERconverge, permulations are applicable to any similar methods. When using permulations to calculate empirical p-values using Forward Genomics, an alternative evolutionary rates-based method, we show that we can quantify a realistic confidence level at which we believe a gene is under accelerated evolution in a subset of species. Even when using the Forward Genomics global method, a deprecated method that does not account for phylogenetic relationships among species, permulations improved the ability to detect accelerated evolution in marine pseudogenes (**Figure 7B**). The improvement is likely due to permulations indirectly capturing phylogenetic information through their construction. For the Forward Genomics local method, permulations captured realistic confidence levels without losing the ability to detect accelerated evolution in marine pseudogenes (**Figures 7B** and **C**). Theoretical p-values directly from the Forward Genomics method (**Figure 7A**) show over half of the genome under significantly accelerated evolution related to the marine phenotype (12,438 out of 18,797 genes with the lowest possible Benjamini Hocberg corrected p-value), which is biologically highly unlikely (Eyre-Walker and Keightley 1999; Eyre-Walker et al. 2002; Eyre-Walker et al. 2006; Kryukov et al. 2007). Permulations reduce the number of genes under significantly accelerated evolutionary rates to a more modest number (889 genes if using the same confidence level cut-off) to more accurately reflect both the biology of the system and our confidence in identifying genes with significant evolutionary rate shifts.

Finally, permulations demonstrate a crucial correction to pathway enrichment statistics that corrects for coevolution among genes in a pathway of interest. Since pathways often contain functionally related genes that evolve at similar rates, performing pathway enrichment treating each gene as an independent observation is statistically incorrect and will result in erroneous conclusions. Performing permulations at the pathway level identifies pathways that are falsely shown to be enriched and correctly quantifies the confidence at which we may state that a pathway is enriched. We argue that a strategy like permulations is essential in virtually all cases of pathway enrichment calculations to account for gene non-independence driven by correlated evolutionary trends.

Overall, permulations are an important statistical consideration that should be undertaken to accurately report results from evolutionary rates-based analyses as presented here. Regardless of whether permulation p-values allow for greater or fewer null hypothesis rejections at a given threshold, they are an accurate depiction of statistical power given a data structure. In the absence of a known parametric null that accurately represents a data set, a permulation-style approach is an important tool to calculate statistical confidence.

## Materials and Methods

### RERconverge

RERconverge finds associations between genetic elements and phenotypes by detecting convergent evolutionary rate shifts in species with convergent phenotypes. The method operates on any type of genetic element and has been used successfully for both protein-coding and noncoding regions. Prior to running RERconverge, phylogenetic trees for each genetic element are generated using the Phylogenetic Analysis by Maximum Likelihood (PAML) program (Yang 2007) or related method, with branch lengths that represent the number of substitutions that occurred between a species and its ancestor. Raw evolutionary rates are converted to relative evolutionary rates (RERs) using RERconverge functions *readTrees* and *getAllResiduals*, which normalize branches for average evolutionary rate along that branch genome-wide and correct for the mean-variance relationship among branch lengths (Partha et al. 2019). RERs and phenotype information are then supplied to *correlateWithBinaryPhenotype* or *correlateWithContinuousPhenotype* functions to calculate element-phenotype associations. Kendall’s Tau associations are calculated for binary phenotypes, and Pearson correlation values are calculated for continuous phenotypes, both by default.

After calculating association statistics, signed log p-values for associations are used to calculate pathway enrichment using the rank-based Wilcoxon Rank-Sum test. The *fastWilcoxGMTAll* function in RERconverge calculates pathway enrichment statistics over a list of pathway annotations using all genes in a particular annotation set as the background.

### Alternate Methods

In addition to performing permulations using RERconverge, we attempted to test our method using PhyloAcc, HyPhy RELAX, and Forward Genomics. Unfortunately, it was not feasible to perform permulation analyses for PhyloAcc and HyPhy RELAX because they were prohibitively computationally expensive. They would require tens of millions of computational hours to generate 500 permulations, the minimum number to accurately represent the null distribution. Therefore, we applied permulations only to the Forward Genomics method to demonstrate the broader applicability of permulations.

Similar to RERconverge, Forward Genomics tests for an accelerated evolutionary rate in foreground branches. The method correlates a normalized substitution rate with the phenotypes using Pearson correlation. Forward Genomics works only for binary phenotypes and has demonstrated success in coding and non-coding elements (Hiller et al. 2012). We used the most recent version of Forward Genomics’ global method, which identifies elements in which all phenotype loss species that have a normalized substitution rate that is higher than in all phenotype-preserving species, and the most recent version of Forward Genomics’ local method, which uses percent identity values per branch to calculate gene-phenotype associations with respect to the underlying phylogeny.

### Phylogenetic Simulations

As shown in **Figure 2**, each permulated phenotype is generated by first performing a phylogenetic simulation using an established phylogenetic topology. To generate the master tree, whose branch lengths represent the average evolutionary rates of all genetic elements in the dataset for each species, the function *readTrees* in RERconverge can be used. Next, the master tree and the trait values (binary or continuous) are used to compute the expected variance of the phenotype per unit time, and subsequently perform a Brownian motion simulation to simulate branch lengths; the R package *GEIGER* (Harmon et al. 2008) is used to perform both operations. Simulated values are then used in different ways for binary and continuous phenotypes to generate permulated phenotypes.

### Details on Binary Permulation

In RERconverge, CC and SSM permulations are performed using the *getPermsBinary* function, by setting the argument “permmode” to “cc” or “ssm”, respectively. The function requires the user to supply information on the original foreground species and their relationships by specifying 1) the names of the extant (tip) foreground species and 2) an R list object containing pair(s) of sister species whose common ancestor(s) is to be included in the foreground set as well (see examples in **Supplementary Walkthrough**). Using these inputs, the function infers the original phenotype tree and assigns the phenotype values to the correct branches (1 for foreground, 0 for background), which is subsequently used as constraints for the permulation. Phylogenetic simulations are then run using the master tree to assign simulated branch lengths to the tree branches.

For the CC permulation, the *n* tip branches with the highest trait values from the simulation, where *n* is the number of observed tip foregrounds, are selected as the new foregrounds. The function then calls the *foreground2Tree* function in RERconverge with “clade” set to “all” to construct a binary tree with a foreground set that includes all branches (tip and internal) in the foreground clades. This simulation is repeated until the number of foregrounds and the phylogenetic relationships among the foregrounds are the same in the observed phenotype and the simulated phenotype.

The SSM permulation matches the tree topology of the permulated phenotypes to the tree of individual genes. To do this, the SSM permulation follows the same steps as described above, with an additional step of trimming off branches that are missing in the gene tree. In this case, the *m* longest tip branches (where *m* is the number of observed tip foregrounds in the *gene* tree) are chosen as new tip foregrounds to run *foreground2Tree*. Thus, in the SSM method, genes with different tree topologies will have different sets of permulations. However, for each unique topology, the number and phylogenetic relationships of the foregrounds are preserved. **Figure 4** shows examples of CC- and SSM-permulated trees for 4 genes with distinct topologies.

### Permulation p-values

After calculating empirical null statistics and p-values, permulation p-values per gene are calculated by finding the proportion of null statistics from permulated phenotypes that are as extreme or more extreme than the statistic calculated using the real phenotype. This proportion represents the proportion of times that we observe a concordance between gene and phenotype evolution as strong as we observed due to random chance given the underlying structure of the data. In RERconverge, the *permpvalcor* function calculates the permulation p-values for a given set of permulation association statistics.

Note that since permulation p-values are a proportion of total permulations, the precision of permulation p-values is based on the total number of permulations performed. For example, with 1,000 permulations, the lowest reportable p-value is 0.001 and permulation p-values calculated as 0 must be reported as <0.001 because we only have precision to report p-values to the thousandths place.

### Permulation p-values for Pathway Enrichment

Empirical null statistics and p-values for pathways are calculated using the empirical null statistics and p-values for individual genes. For each set of empirical null statistics generated from a particular permulated phenotype, genes are assigned the log of the empirical null p-value times the sign of the empirical null statistic for that permulation. Empirical null pathway statistics are calculated for each permulation using those values with the RERconverge function *fastWilcoxGMTall* that performs a Wilcoxon Rank-Sum test comparing values from genes in a pathway to values in background genes. The function *getEnrichPerms* calculates null enrichment statistics given a set of null correlation statistics, or, alternatively, *getPermsBinary* and *getPermsContinuous* calculate both null correlation and null pathway enrichment statistics simultaneously by default for the binary and continuous phenotypes, respectively. Permulation p-values for pathway enrichment are then calculated as the proportion of empirical null statistics that are as extreme or more extreme than the observed enrichment statistic using the *permpvalenrich* function. Pathways that show significant parametric p-values and non-significant permulation p-values are likely cases of genes “moving in packs” and are not truly significantly enriched.

## Supporting information

Supplementary Walkthrough

## Data Availability Statement

The data underlying this article are available in the RERconverge repository on github (https://github.com/nclark-lab/RERconverge). The data for the long-lived large-bodied phenotype are publicly available on Anage (https://genomics.senescence.info/species/index.html), and has been previously published in (Kowalczyk et al. 2020).

## Acknowledgement

This work was supported by the National Institutes of Health (R01 HG009299 to N.C. and M.C and T32 EB009403 to A.K.).

## References

Allison DB, Gadbury GL, Heo M, Fernández JR, Lee C-K, Prolla TA, Weindruch R. 2002. A mixture model approach for the analysis of microarray gene expression data. Computational Statistics & Data Analysis. 39(1):1–20. doi:10.1016/S0167-9473(01)00046-9.

Bininda-Emonds ORP, Cardillo M, Jones KE, MacPhee RDE, Beck RMD, Grenyer R, Price SA, Vos RA, Gittleman JL, Purvis A. 2007. The delayed rise of present-day mammals. Nature. 446(7135):507–512. doi:10.1038/nature05634.

Boyle EI, Weng S, Gollub J, Jin H, Botstein D, Cherry JM, Sherlock G. 2004. GO::TermFinder--open source software for accessing Gene Ontology information and finding significantly enriched Gene Ontology terms associated with a list of genes. Bioinformatics. 20(18):3710–3715. doi: 10.1093/bioinformatics/bth456.

Chikina M, Robinson JD, Clark NL. 2016. Hundreds of Genes Experienced Convergent Shifts in Selective Pressure in Marine Mammals. Mol Biol Evol. 33(9):2182–2192. doi: 10.1093/molbev/msw112.

Clark NL, Alani E, Aquadro CF. 2012. Evolutionary rate covariation reveals shared functionality and coexpression of genes. Genome Res. 22(4):714–720. doi:10.1101/gr.132647.111.

Clark NL, Alani E, Aquadro CF. 2013. Evolutionary rate covariation in meiotic proteins results from fluctuating evolutionary pressure in yeasts and mammals. Genetics. 193(2):529–538. doi:10.1534/genetics.112.145979.

Eden E, Lipson D, Yogev S, Yakhini Z. 2007. Discovering Motifs in Ranked Lists of DNA Sequences. Fraenkel E, editor. PLoS Comput Biol. 3(3):e39. doi:10.1371/journal.pcbi.0030039.

Eden E, Navon R, Steinfeld I, Lipson D, Yakhini Z. 2009. GOrilla: a tool for discovery and visualization of enriched GO terms in ranked gene lists. BMC Bioinformatics. 10(1):48. doi:10.1186/1471-2105-10-48.

Eyre-Walker A, Keightley PD. 1999. High genomic deleterious mutation rates in hominids. Nature. 397(6717):344–347. doi:10.1038/16915.

Eyre-Walker A, Keightley PD, Smith NGC, Gaffney D. 2002. Quantifying the Slightly Deleterious Mutation Model of Molecular Evolution. Molecular Biology and Evolution. 19(12):2142–2149. doi: 10.1093/oxfordjournals.molbev.a004039.

Eyre-Walker A, Woolfit M, Phelps T. 2006. The Distribution of Fitness Effects of New Deleterious Amino Acid Mutations in Humans. Genetics. 173(2):891–900. doi: 10.1534/genetics.106.057570.

Felsenstein J. 1985. Phylogenies and the Comparative Method. The American Naturalist. 125(1): 1–15.

Fraser HB, Hirsh AE, Wall DP, Eisen MB. 2004. Coevolution of gene expression among interacting proteins. PNAS. 101(24):9033–9038. doi:10.1073/pnas.0402591101.

Harmon LJ, Weir JT, Brock CD, Glor RE, Challenger W. 2008. GEIGER: investigating evolutionary radiations. Bioinformatics. 24(1):129–131. doi:10.1093/bioinformatics/btm538.

Hiller M, Schaar BT, Indjeian VB, Kingsley DM, Hagey LR, Bejerano G. 2012. A “forward genomics” approach links genotype to phenotype using independent phenotypic losses among related species. Cell Rep. 2(4):817–823. doi:10.1016/j.celrep.2012.08.032.

Hu X, Gadbury GL, Xiang Q, Allison DB. 2010. Illustrations on Using the Distribution of a P-value in High Dimensional Data Analyses. Adv Appl Stat Sci. 1(2): 191–213.

Hu Z, Sackton TB, Edwards SV, Liu JS. 2019. Bayesian Detection of Convergent Rate Changes of Conserved Noncoding Elements on Phylogenetic Trees. Pond SK, editor. Molecular Biology and Evolution. 36(5):1086–1100. doi:10.1093/molbev/msz049.

Juan D, Pazos F, Valencia A. 2008. High-confidence prediction of global interactomes based on genome-wide coevolutionary networks. Proceedings of the National Academy of Sciences. 105(3):934–939. doi: 10.1073/pnas.0709671105.

Kowalczyk A, Meyer WK, Partha R, Mao W, Clark NL, Chikina M. 2019. RERconverge: an R package for associating evolutionary rates with convergent traits. Valencia A, editor. Bioinformatics. 35(22):4815–4817. doi:10.1093/bioinformatics/btz468.

Kowalczyk A, Partha R, Clark NL, Chikina M. 2020. Pan-mammalian analysis of molecular constraints underlying extended lifespan. eLife. 9:e51089. doi:10.7554/eLife.51089.

Kryukov GV, Pennacchio LA, Sunyaev SR. 2007. Most Rare Missense Alleles Are Deleterious in Humans: Implications for Complex Disease and Association Studies. The American Journal of Human Genetics. 80(4):727–739. doi:10.1086/513473.

Majewski IJ, Ritchie ME, Phipson B, Corbin J, Pakusch M, Ebert A, Busslinger M, Koseki H, Hu Y, Smyth GK, et al. 2010. Opposing roles of polycomb repressive complexes in hematopoietic stem and progenitor cells. Blood. 116(5):731–739. doi:10.1182/blood-2009-12-260760.

Meredith RW, Janecka JE, Gatesy J, Ryder OA, Fisher CA, Teeling EC, Goodbla A, Eizirik E, Simao TLL, Stadler T, et al. 2011. Impacts of the Cretaceous Terrestrial Revolution and KPg Extinction on Mammal Diversification. Science. 334(6055):521–524. doi:10.1126/science.1211028.

Meyer WK, Jamison J, Richter R, Woods SE, Partha R, Kowalczyk A, Kronk C, Chikina M, Bonde RK, Crocker DE, et al. 2018. Ancient convergent losses of *Paraoxonase 1* yield potential risks for modern marine mammals. Science. 361(6402):591–594. doi:10.1126/science.aap7714.

Nettleton D, Hwang JTG, Caldo RA, Wise RP. 2006. Estimating the number of true null hypotheses from a histogram of p values. JABES. 11(3):337–356. doi:10.1198/108571106X129135.

Pagel M, Meade A. 2006. Bayesian Analysis of Correlated Evolution of Discrete Characters by Reversible-Jump Markov Chain Monte Carlo. The American Naturalist. 167(6):808–825. doi:10.1086/503444.

Partha R, Chauhan BK, Ferreira Z, Robinson JD, Lathrop K, Nischal KK, Chikina M, Clark NL. 2017. Subterranean mammals show convergent regression in ocular genes and enhancers, along with adaptation to tunneling. eLife. 6:e25884. doi:10.7554/eLife.25884.

Partha R, Kowalczyk A, Clark NL, Chikina M. 2019. Robust Method for Detecting Convergent Shifts in Evolutionary Rates. Tamura K, editor. Molecular Biology and Evolution. 36(8):1817–1830. doi:10.1093/molbev/msz107.

Prudent X, Parra G, Schwede P, Roscito JG, Hiller M. 2016. Controlling for Phylogenetic Relatedness and Evolutionary Rates Improves the Discovery of Associations Between Species’ Phenotypic and Genomic Differences. Mol Biol Evol. 33(8):2135–2150. doi:10.1093/molbev/msw098.

Ritchie ME, Phipson B, Wu D, Hu Y, Law CW, Shi W, Smyth GK. 2015. limma powers differential expression analyses for RNA-sequencing and microarray studies. Nucleic Acids Res. 43(7):e47. doi: 10.1093/nar/gkv007.

Stone GN, Nee S, Felsenstein J. 2011. Controlling for non-independence in comparative analysis of patterns across populations within species. Philos Trans R Soc Lond, B, Biol Sci. 366(1569):1410–1424. doi:10.1098/rstb.2010.0311.

Storey JD, Bass AJ, Dabney A, Robinson D. 2020. qvalue: Q-value estimation for false discovery rate control. http://github.com/jdstorey/qvalue.

Storey JD, Tibshirani R. 2003. Statistical significance for genomewide studies. Proceedings of the National Academy of Sciences. 100(16):9440–9445. doi:10.1073/pnas.1530509100.

Subramanian A, Tamayo P, Mootha VK, Mukherjee S, Ebert BL, Gillette MA, Paulovich A, Pomeroy SL, Golub TR, Lander ES, et al. 2005. Gene set enrichment analysis: A knowledge-based approach for interpreting genome-wide expression profiles. Proceedings of the National Academy of Sciences. 102(43):15545–15550. doi:10.1073/pnas.0506580102.

Wertheim JO, Murrell B, Smith MD, Kosakovsky Pond SL, Scheffler K. 2015. RELAX: Detecting Relaxed Selection in a Phylogenetic Framework. Molecular Biology and Evolution. 32(3):820–832. doi: 10.1093/molbev/msu400.

Wu D, Smyth GK. 2012. Camera: a competitive gene set test accounting for inter-gene correlation. Nucleic Acids Research. 40(17):e133–e133. doi:10.1093/nar/gks461.

Yang Z. 2007. PAML 4: Phylogenetic Analysis by Maximum Likelihood. Molecular Biology and Evolution. 24(8):1586–1591. doi:10.1093/molbev/msm088.

