## Supplementary Walkthrough for "Phylogenetic Permulations: a statistically rigorous approach to measure confidence in associations between phenotypes and genetic elements in a phylogenetic context"

### Permulation Walkthrough

This walkthrough provides instructions to perform permulation analysis to calculate empirical p-values for genes and pathways. Due to a non-uniform p-value distribution for gene-evolutionary rate associations and non-independence among genes for pathway enrichment, parametric p-values directly from RERconverge may not accurately represent true confidence of association or enrichment. Permulation p-values correct this issue. For the continuous phenotype, parametric p-values for gene correlations tend to be overestimated and parametric p-values for pathway enrichment tend to be underestimated.

Permulations are performed using the following steps after performing standard RERconverge analyses:

1. generate null (permulated) phenotypes
2. recalculate gene correlation and pathway enrichment statistics using null phenotypes
3. quantify the proportion of null statistics more significant than observed statistics

Since statistics generated from null phenotypes represent the true null distribution for the statistic calculated over the given data, the proportion of extreme null statistics quantifies true confidence in the extremity of a particular correlation or enrichment. These are permulation p-values and can be interpreted similarly to standard p-values, including performing multiple hypothesis testing correction. One caveat is that the precision of a permulation p-value is limited by the number of permulations performed - the lowest observable p-value is the reciprocal of the number of permulations performed.

Permulations are a combination of phylogenetic simulations and permutations. To generate a permulated phenotype, first a simulated phenotype is generated based on a phylogeny with branch lengths that represent the average genome-wide evolutionary rate along that branch using a Brownian motion model of evolution. Observed phenotype values are then assigned to species based on the ranks of simulated values - the highest simulated value is assigned the highest observed value, the second-highest simulated values is assigned the second-highest observed value, etc. Permulations are favored over permulations because they respect the underlying phylogenetic relationships among species, so more closely related species have more similar phenotypes. Permulations are favored over simulations because they preserve the exact distribution and range of the observed phenotype. In practice, permulation p-values are more conservative (i.e. lower) than permutation p-values and equally as conservative as simulation p-values.

**In R, permulation procedures are conveniently bundled into a handful of functions for simplicity. Note that these functions can take a very long time to run on large datasets and for large numbers of permulations.**

#### Binary Permulations

To evaluate the association between relative evolutionary rates (RERs) and **binary traits**, we first have to provide a binary trait tree that specifies which branches in the tree have the trait of interest (**foreground branches**). In this vignette, we'll use the marine mammals phenotype to demonstrate the steps. Let's first read the set of trees for all the genes with the `readTrees` function.

```
library(RERconverge)
rerpath = find.package('RERconverge')
```

```
#read trees
toytreefile = "subsetMammalGeneTrees.txt"
toyTrees=readTrees(paste(rerpath,"/extdata/",toytreefile,sep=""), max.read = 200)
```

The foreground set for the marine mammals phenotype include 3 independent tip branches (walrus, seal, manatee) and the cetacean clade (killer whale, dolphin, and the cetacean ancestor). Walrus and seal branched out from a common ancestor, but the common ancestor is not included in the foreground set. To set up a foreground set that contains a mixture of independent tip species and clades, we can use the `foreground2TreeClades` function. This function requires the following input:

- `fg_vec`: a vector containing the names of the tip foreground species
- `sisters_list`: a list object containing information on clades in the foreground set, specifically the pair(s) of "sister species" that branch out from the same ancestor
- `trees`: trees object from `readTrees`

For example, the marine foregrounds can be constructed as follows:

```
marineFg = c("Killer_whale", "Dolphin", "Walrus", "Seal", "Manatee")
sisters_marine = list("clade1"=c("Killer_whale", "Dolphin"))

marineFgTree = foreground2TreeClades(marineFg,sisters_marine,toyTrees,plotTree=F)
marineplot1 = plotTreeHighlightBranches(marineFgTree,
                                        hlspecies=which(marineFgTree$edge.length==1),
                                        hlcols="blue", main="Marine mammals trait tree")
```

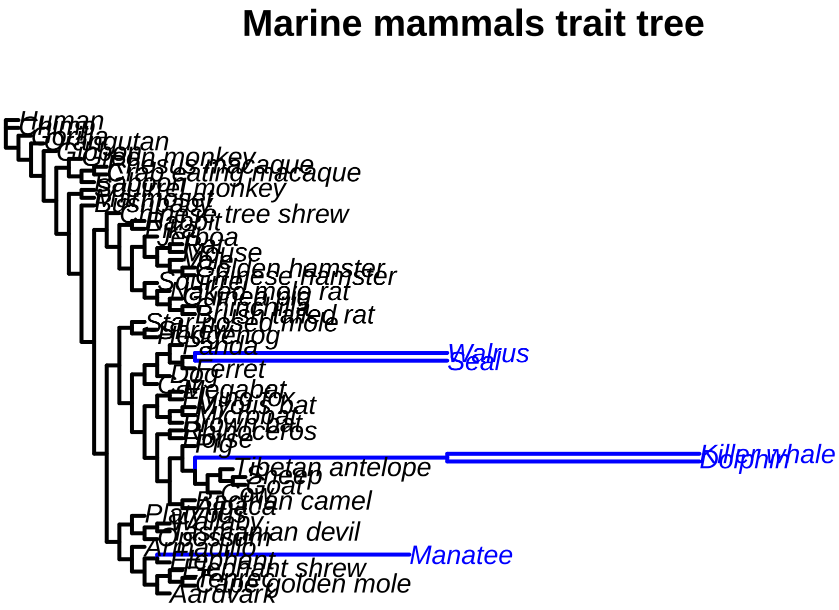


Notice that in the resulting binary trait tree, the ancestor of walrus and seal are not included in the foreground set. If this ancestor should also be included, the `sisters_list` input should be specified as follows:

```
sisters_marine2 = list("clade1"=c("Killer_whale","Dolphin"), "clade2"=c("Walrus","Seal"))
marineFgTree2 = foreground2TreeClades(marineFg,sisters_marine2,toyTrees,plotTree=F)
marineplot2 = plotTreeHighlightBranches(marineFgTree2,
                    hlspecies=which(marineFgTree2$edge.length==1),
                    hlcols="blue",
                    main="Modified marine trait tree with walrus-seal ancestor")
```

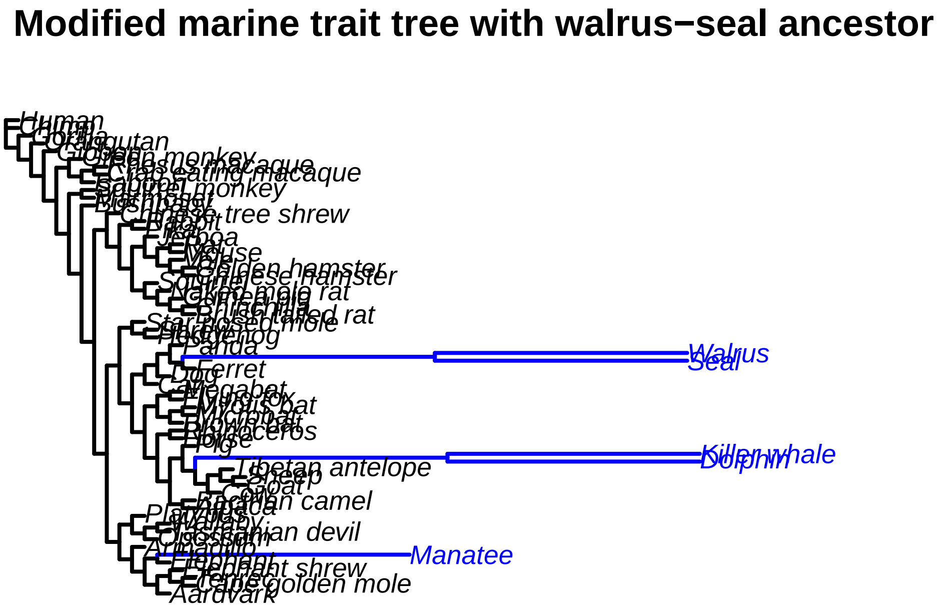


Thus, the `foreground2TreeClades` function allows for flexible definition of foregrounds that include tip branches and clades with any depth. Below is an example of how to specify foreground clades where the ancestor of an ancestor is also a foreground species:

```
fgExample = c("Golden_hamster","Chinese_hamster","Vole","Walrus","Seal")
sisters_Example = list("clade1"=c("Golden_hamster","Chinese_hamster"),
                       "clade2"=c("clade1","Vole"))

exampleFgTree = foreground2TreeClades(fgExample,sisters_Example,toyTrees,plotTree=F)

exampleplot = plotTreeHighlightBranches(exampleFgTree,
                    hlspecies=which(exampleFgTree$edge.length==1),
                    hlcols="blue",
                    main="Example trait tree with whole clade foreground")
```

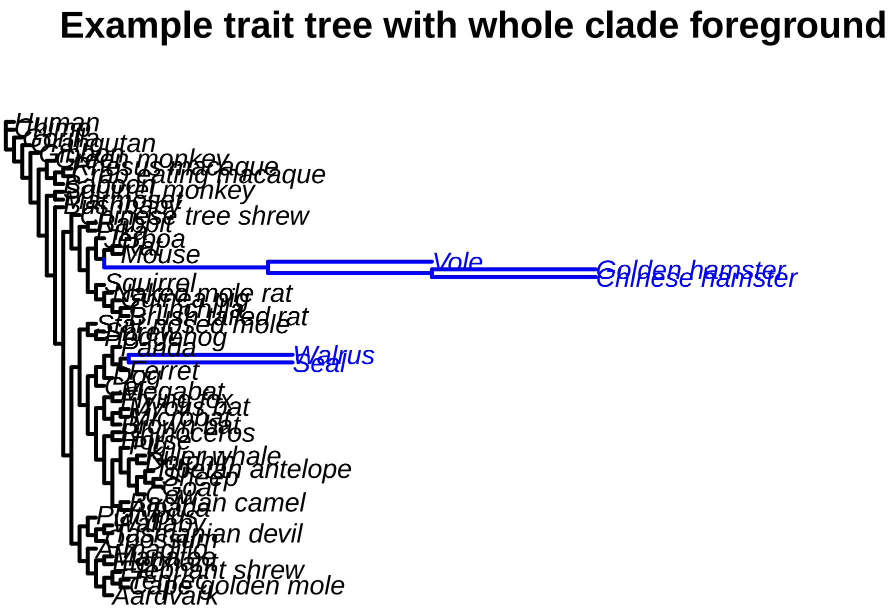


Back to the marine phenotype example, once we have the binary trait tree, we can calculate the actual correlations with RERs following the steps below. Note that when `foreground2TreeClades`is used, the `tree2PathsClades` function should be used to compute the paths, instead of the standard `tree2Paths` like in the Full Walkthrough.

```
# Calculating paths from the foreground tree
pathvec = tree2PathsClades(marineFgTree, toyTrees)

# Calculate RERs
mamRERw = getAllResiduals(toyTrees, transform="sqrt", weighted=T, scale=T)
```

```
# Calculate correlation
res = correlateWithBinaryPhenotype(mamRERw, pathvec, min.sp=10, min.pos=2,
                                    weighted="auto")
```

After calculating the actual correlation statistics, we can proceed with calculating the permulated correlations. The function `getPermsBinary` performs permulations of binary traits and produces the null p-values, correlation statistics, and enrichment by taking in the following input:

- `numperms`: the number of permulations to perform. Note that the total number of permulations is the limit to permulation p-value precision - the lowest possible permulation p-value is 1/numperms
- `fg_vec`: a vector containing the names of tip foreground species
- `sisters_list`: a list object containing information on clades in the foreground set, specifically the pair(s) of "sister species" that branch out from the same ancestor
- `root_sp`: the species to root the trees on
- `RERmat`: RER matrix from `getAllResiduals` as used in `correlateWithBinaryPhenotype`
- `trees`: trees object from `readTrees`
- `mastertree`: rooted and fully dichotomous tree containing all species with branch lengths representing

average evolutionary rate genome wide. In most cases, modify the master tree from `trees`
- `permmode`: default "cc". Set to "cc" to use the Complete Case method, or "ssm" to use the Species Subset Match method.
- `method`: default "k" for the Kendall Tau test (for binary traits)
- `trees_list`: default NULL. If `permmode`="ssm", this (optional) input specifies the list of gene trees to perform permulations for. If `permmode`="ssm" and `trees_list`=NULL, permulations will be performed for all gene trees. Set this input to NULL if `permmode`="cc".
- `calculateenrich`: default F. Boolean specifying if permulation enrichment statistics should be calculated
- `annotlist`: annotations as used in `fastwilcoxGMTall`. Not used if `calculateenrich`=F

To run binary permulations with the Complete Case (CC) method, follow the example in the block of code below, most importantly setting `permmode`="cc". The output of `getPermsBinary`, which contains the p-values and correlation statistics of the permulations, should then be supplied to `permpvalcor` to compute the empirical permulation p-values of the genes from the permulated correlation statistics.

```
#define the root species
root_sp = "Human"

masterTree = toyTrees$masterTree

#perform binary CC permulation
permCC = getPermsBinary(100, marineFg, sisters_marine, root_sp, mamRERw, toyTrees,
                        masterTree, permmode="cc")

#calculate permulation p-values
permpvalCC = permpvalcor(res,permCC)
```

If we want to calculate the enrichment permulation statistics simultaneously, use the same function and set `calculateenrich`=T, as follows:

```
#load annotations
annots=RERconverge::read.gmt("gmtfile.gmt")
annotlist=list(annots)
names(annotlist)="MSigDBpathways"

#perform permulations
permCCWithCor = getPermsBinary(20, marineFg, sisters_marine, root_sp, mamRERw, toyTrees,
                               masterTree, permmode="cc",calculateenrich = T,
                               annotlist=annotlist)
```

To run the Species Subset Match (SSM) permulation, we can use the same function `getPermsBinary` while setting `permmode`="ssm". Note that because the SSM method produces distinct sets of permulations for different genes, this method is much more computationally intensive and takes a significantly longer time compared to the CC method. Hence, we advise that the SSM method should be run in batches of smaller numbers of permulations. The outputs of different batches can then be combined using the `combinePermData` function, such as shown in the example below. Let's perform 10 permulations of the first 10 gene trees in the list.

```
#specify the list of trees to permulate
trees_example = toyTrees$trees[1:10]

#perform 2 batches of binary SSM permulations
permSSM1 = getPermsBinary(5, marineFg, sisters_marine, root_sp, mamRERw, toyTrees,
                          masterTree, permmode="ssm",trees_example)
permSSM2 = getPermsBinary(5, marineFg, sisters_marine, root_sp, mamRERw, toyTrees,
                          masterTree, permmode="ssm",trees_example)
```

```
#combine the outputs of 2 batches
combpermSSM = combinePermData(permSSM1, permSSM2, enrich=F)

#calculate permulation p-values
permpvalSSM = permpvalcor(res,combpermSSM)
```

RERconverge also allows user to produce permulated binary phenotype trees to use with other softwares (e.g., ForwardGenomics, HyPhy RELAX, PhyloAcc, etc.), without calculating permulated correlation statistics. The functions `simBinPhenoCC` and `simBinPhenoSSM` produce one permulated phenotype using the CC and SSM methods, respectively. The code below produces one CC permulation:

```
#producing one permulated tree using CC permulation
treeCC = simBinPhenoCC(toyTrees, masterTree, root_sp, marineFg, sisters_marine, pathvec,
                       plotTreeBool=F)
CCtree = plotTreeHighlightBranches(treeCC,hlspecies=which(treeCC$edge.length==1),
                                   hlcols="blue")
```

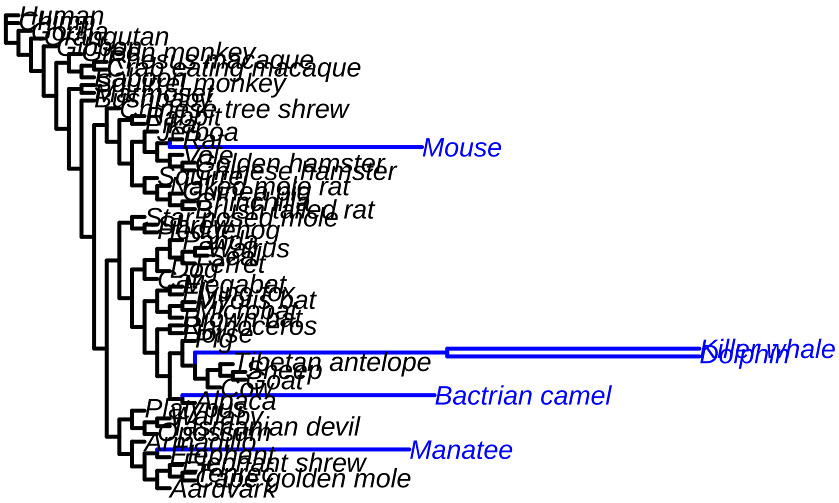


For SSM, the permulated tree of a gene may be different from that of another gene if they have different "species membership", meaning if the gene trees may be missing different sets of species. Compared to `simBinPhenoCC`, `simBinPhenoSSM` requires an additional input `tree`, which specifies the gene tree of the specific gene to permulate. Below is an example permulation of a gene that has all marine foregrounds in its tree:

```
#producing one permulated tree for a gene with all marine foregrounds using SSM
TTNtree = toyTrees$trees$TTN

#plot the gene tree of TTN
```

```
TTNtreePlot = plotTreeHighlightBranches(TTNtree,
                                        hlspecies=which(marineFgTree$edge.length==1),
                                        hlcols="blue",
                                        main="TTN tree") #plot individual gene tree
```

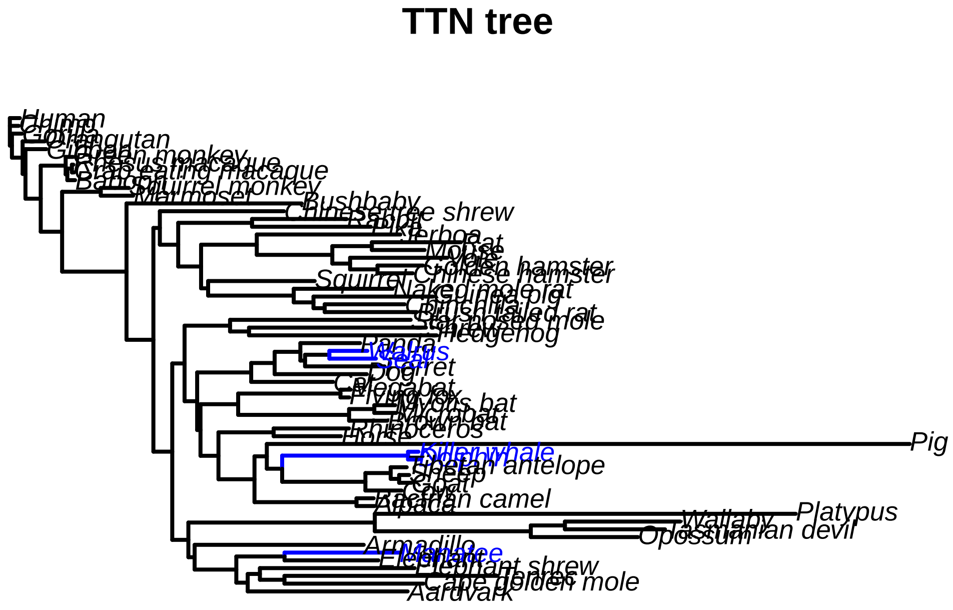


```
#generate and plot a SSM permulation of TTN
TTNtreeSSM = simBinPhenoSSM(TTNtree, toyTrees, root_sp, marineFg, sisters_marine, pathvec,
                            plotTreeBool=F)
TTNplotSSM = plotTreeHighlightBranches(TTNtreeSSM,
                                       hlspecies=which(TTNtreeSSM$edge.length==1),
                                       hlcols="blue")
```

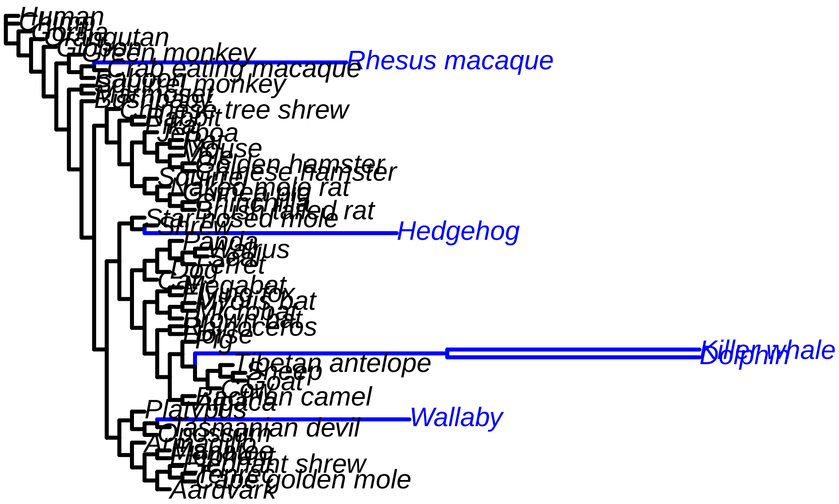


And below is an example permulation of a gene that is missing some marine foregrounds in its tree:

```
#producing one permulated tree for a gene that is missing some marine foregrounds
ACBD5tree = toyTrees$trees$ACBD5
ind.marine = which(ACBD5tree$tip.label %in% marineFg)
marineFgACBD5 = ACBD5tree$tip.label[ind.marine]

#plot the gene tree of ACBD5
ACBD5treePlot = plotTreeHighlightBranches(ACBD5tree,
                                    hlspecies=marineFgACBD5,
                                    hlcols=rep("blue",length(marineFgACBD5)),
                                    main="ACBD5 tree")
```

#### ACBD5 tree

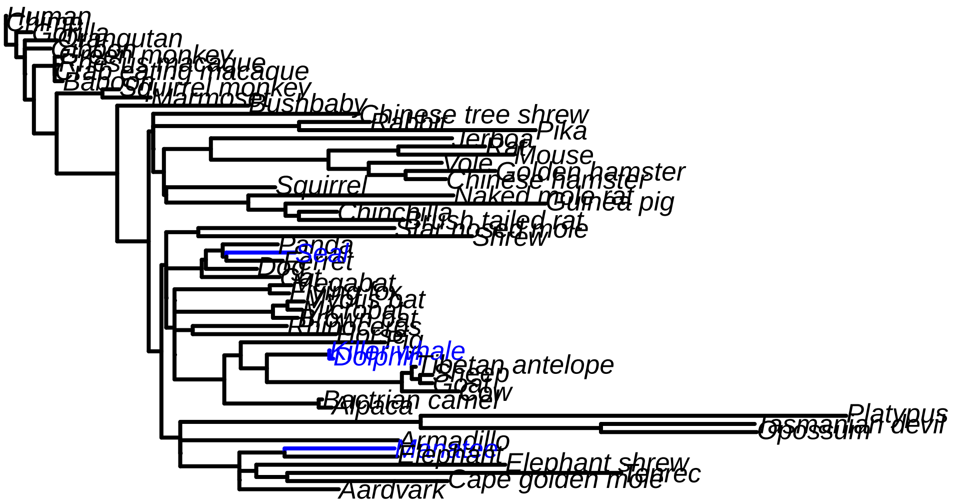


```r
#generate and plot one SSM permulation of ACBD5
ACBD5treeSSM = simBinPhenoSSM(ACBD5tree, toyTrees, root_sp, marineFg, sisters_marine,
                              pathvec, plotTreeBool=F)
ACBD5plotSSM = plotTreeHighlightBranches(ACBD5treeSSM,
                                         hlspecies=which(ACBD5treeSSM$edge.length==1),
                                         hlcols="blue")
```

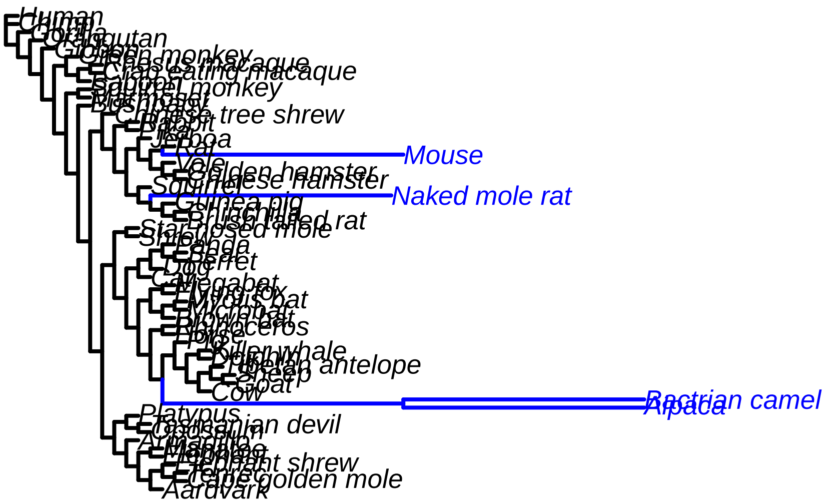


To produce multiple permulations using the CC method, the function `generatePermulatedBinPhen` can be used, specifying `permmode`="cc" and supplying the master tree to the input `tree`. For example, the code below produces 20 CC permulated trees:

```
# number of permulations
numperms = 20

TTNpermulatedTreesCC = generatePermulatedBinPhen(masterTree, numperms, toyTrees, root_sp,
                                                 marineFg, sisters_marine, pathvec,
                                                 permmode="cc")
```

To produce multiple permulations of ONE gene using the SSM method, `generatePermulatedBinPhen` can also be used with `permmode`="ssm". The code below produces 20 SSM permulations of TTN:

```
# number of permulations
numperms = 20

TTNpermulatedTreesSSM = generatePermulatedBinPhen(TTNtree, numperms, toyTrees, root_sp,
                                                  marineFg, sisters_marine, pathvec,
                                                  permmode="ssm")
```

RERconverge also allows users to generate SSM permulations for **multiple** genes at the same time. However, since the SSM method is a lot more computationally intensive compared to the CC method, permulations are run in 'batches' based on distinct species membership. To do this, use the function `generatePermulatedBinPhenSSMBatched`, which requires the following input:

- `trees_list`: the list of gene trees to produce permulations for (e.g., a subset of toyTrees$trees)
- `numperms`: integer number of permulations

- `trees`: trees object from `readTrees`
- `root_sp`: the species to root the trees on
- `fg_vec`: a vector of names of tip foreground species
- `sisters_list`: a list object containing information on clades in the foreground set, specifically the pair(s) of "sister species" that branch out from the same ancestor
- `pathvec`: a path vector calculated by `foreground2Paths`, `tree2Paths`, or `tree2PathsClades`

For example, the code below generates 5 permulations of the first 10 trees in the list:

```
# list of gene trees
trees_example = toyTrees$trees[1:10]

# number of permulations
numperms = 5

SSMpermulatedTreesBatched = generatePermulatedBinPhenSSMBatched(trees_example, numperms,
                                toyTrees, root_sp, marineFg, sisters_marine, pathvec)
```

#### Continuous Permulations

First, conduct standard RERconverge analysis. Please see full walkthroughs for more details about these steps.

```
#load RER package
#library(RERconverge)
#rerpath = find.package('RERconverge')

#read trees
toytreefile = "subsetMammalGeneTrees.txt"
toyTrees=RERconverge::readTrees(paste(rerpath,"/extdata/",toytreefile,sep=""),
                                max.read = 200)

#load phenotype data
data("logAdultWeightcm")

#calculate RERs
mamRERw = RERconverge::getAllResiduals(toyTrees,useSpecies=names(logAdultWeightcm),
                                       transform = "sqrt", weighted = T, scale = T)
```

```
#generate trait tree
charpaths=RERconverge::char2Paths(logAdultWeightcm, toyTrees)

#calculate correlation statistics
res=RERconverge::correlateWithContinuousPhenotype(mamRERw, charpaths, min.sp = 10,
                                                   winsorizeRER = 3, winsorizetrait = 3)

#calculate pathway enrichment statistics
stats=RERconverge::getStat(res)
annots=RERconverge::read.gmt("gmtfile.gmt")
annotlist=list(annots)
names(annotlist)="MSigDBpathways"
enrichment=RERconverge::fastwilcoxGMTall(stats, annotlist, outputGeneVals=T, num.g=10)
```

After calculating parametric statistics above, calculate permulations. The `getPermsContinuous` function operates by generating null p-values and statistics for gene correlations and enrichment statistics. The function requires the following input:

- `numperms`: the number of permulations to perform, recommended at least 1000. Note that the total number of permulations is the limit to permulation p-value precion - the lowest possible permulation p-value is 1/numperms
- `traitvec`: phenotype vector as specified in `correlateWithContinuousPhenotype`
- `RERmat`: RER matrix from `getAllResiduals` as used in `correlateWithContinuousPhenotype`
- `annotlist`: annotations as used in `fastwilcoxGMTall`. Not used if `calculateenrich`=F
- `trees`: trees object from `readTrees` as used in `getAllResiduals`
- `mastertree`: rooted and fully dichotomous tree containing all species with branch lengths representing average evolutionary rate genome wide. In most cases, modify the master tree from `trees`
- `calculateenrich`: default T. Boolean specifying if permulation enrichment statistics should be calculated
- `type`: default "simperm". Specifies method to generate null phenotypes. "simperm" specifies permulations, "sim" specifies phylogenetic simulations, and "perm" specifies permutations
- `winR` and `winT`: default 3. Numeric values specifying how much to winsorize RER and trait trees, respectively. Should match `winR` and `winT` values used in `correlateWithContinuousPhenotype`

**This example uses only 100 permulations as a toy example. In practice, we have found that for this group of species and this gene set, at least 500 permulations should be performed to obtain meaningful p-values. Ideally, as many permulations as possible should be performed to maximize p-value precision.**

```
mt=toyTrees$masterTree
mt=root.phylo(mt, outgroup="Platypus", resolve.root=T)

perms=RERconverge::getPermsContinuous(100, logAdultWeightcm, mamRERw, annotlist,
                                      toyTrees, mt)
corpermpvals=RERconverge::permpvalcor(res, perms)
enrichpermpvals=RERconverge::permpvalenrich(enrichment, perms)

# add permulations to real results

res$permpval=corpermpvals[match(rownames(res), names(corpermpvals))]
res$permpvaladj=p.adjust(res$permpval, method="BH")
count=1
while(count<=length(enrichment)){
  enrichment[[count]]$permpval=enrichpermpvals[[count]][match(rownames(enrichment[[count]]),
                                                  names(enrichpermpvals[[count]]))]
  enrichment[[count]]$permpvaladj=p.adjust(enrichment[[count]]$permpval, method="BH")
  count=count+1
}
```

As an alternative to running correlation and enrichment analyses simultaneously, the `getPermsContinuous` function may be used to run correlation analyses alone, and then the `getEnrichPermsContinuous` function may be used to calculate null enrichment statistics based on the null correlation statistics. In this case, the output from `getEnrichPermsContinuous` should be supplied to `permpvalcor` and `permpvalenrich` functions.

This pipeline may be useful in the following cases 1. If very large datasets with many datasets are batched to run `readTrees`, `getAllResiduals`, and to calculate gene correlation statistics and then combined during standard RERconverge analysis, the following procedure should be used to mimic those steps during permulation analyses 2. If new pathway annotations are being tested after gene correlation permulations have already been run, `getEnrichPermsContinuous` can be used to calculate permulation enrichment statistics without rerunning correlation analyses.

```r
permsnoenrich=RERconverge::getPermsContinuous(100, logAdultWeightcm, mamRERw, annotlist,
                                              toyTrees, mt, calculateenrich = F)
permswithenrich=RERconverge::getEnrichPermsContinuous(permsnoenrich, enrichment,
                                                      annotlist)
corpermpvals2=RERconverge::permpvalcor(res, permswithenrich)
enrichpermpvals2=RERconverge::permpvalenrich(enrichment, permswithenrich)

plot(corpermpvals,corpermpvals2)
```

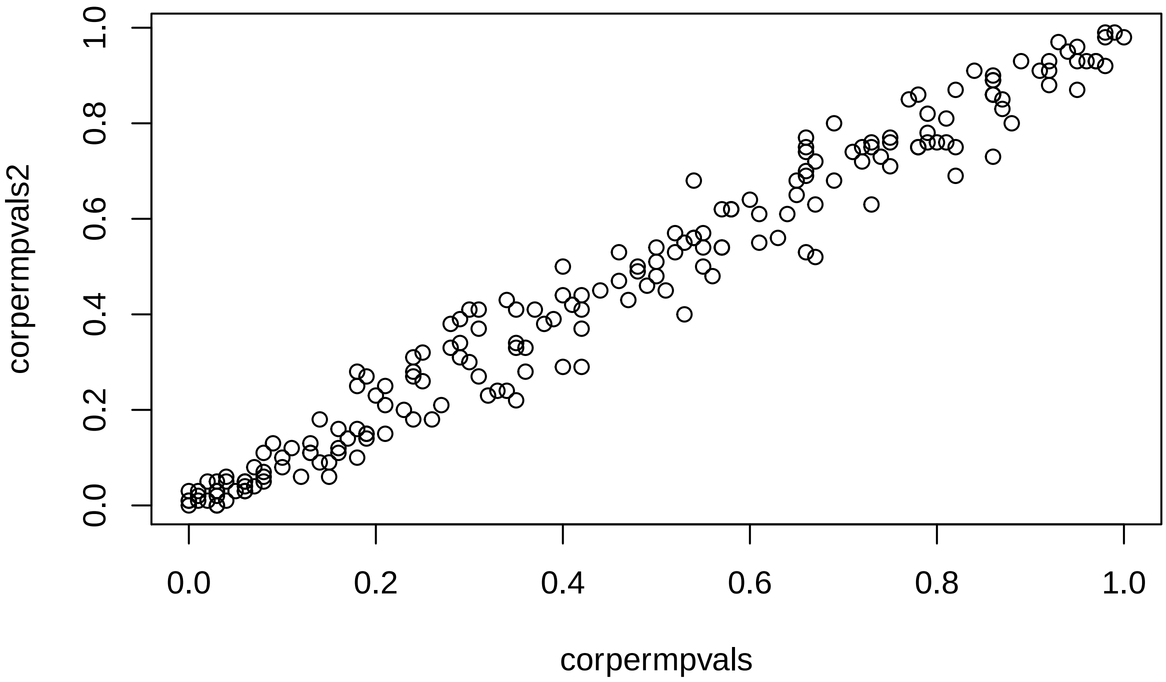


```r
plot(enrichpermpvals[[1]],enrichpermpvals2[[1]])
```

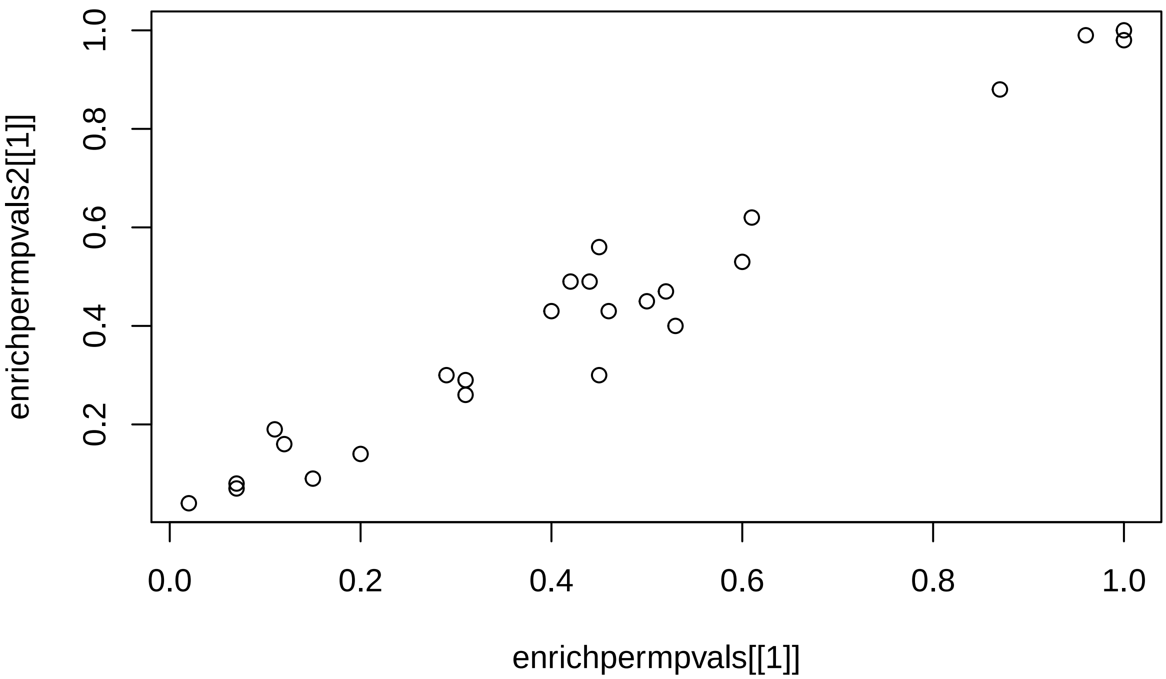


Note variations in these two separate sets of permulation p-values. This stochasticity highlights the necessity of running many permulations to explore as much null phenotype space as possible.

Permulations can also be run in batches and combined using `combinePermData`. This function would be useful for combining several permulation batches run in tandem for computational efficiency.

```
perm2=RERconverge::getPermsContinuous(100, logAdultWeightcm, mamRERw, annotlist,
                                      toyTrees, mt)
combperms=RERconverge::combinePermData(perms, perm2)
```

Subsequent analyses should include subsetting observed significant pathways and genes according to parametric statistics based on significance according to permulation statistics.
